# Year-round rhythms: alpine plant species modulate soil and microbial dynamics during the growing season and under the snow

**DOI:** 10.1101/2025.02.18.638831

**Authors:** Adam Taylor Ruka, Johannes Schweichhart, Jiri Dolezal, Kateřina Čapková, Zuzana Chlumská, Rosa Paulina Calvillo-Medina, Travis B. Meador, Nadine Praeg, Paul Illmer, Roey Angel, Klara Rehakova

**Affiliations:** Institute of Botany CAS, Třeboň, Czechia; Institute of Soil Biology and Biogeochemistry, Biology Centre CAS, České Budějovice, Czechia; Faculty of Science, University of South Bohemia in České Budějovice, České Budějovice, Czechia; Institute of Hydrobiology, Biology Centre CAS, České Budějovice, Czechia; Department of Microbiology, Universität Innsbruck, Innsbruck, Austria

**Keywords:** Seasonality, Phenology, Microbial, Winter, Snow, Biogeochemical, Alpine

## Abstract

1. Soil-plant-microbe interactions are integral throughout most terrestrial ecosystems, yet the importance of plant phenology and seasonal dynamism upon these relationships remains unknown. Given the pronounced seasonality of alpine environments, we sampled 8 plant species occurring in two habitats (alpine meadow and subnival zone) across four seasons (including snow-covered winter) in the Central Eastern Alps to determine the plant growth strategies and plant nutrient parameters which closely couple with rhizosphere microbial parameters.
2. In subnival locations, plants exhibited stronger seasonal changes among leaf and root tissue nutrient concentrations and non-structural carbohydrates (NSCs) compared to those in lower elevation alpine meadows. However, rhizosphere microbial parameters (microbial biomass (MBC), extracellular enzymes, and community composition) demonstrated more seasonal changes in the alpine meadow locations.
3. A phenological delay was observed in bacterial and fungal communities of the subnival zone, with peak plant rhizosphere differentiation occurring later in the season than in alpine meadows. Therefore, the prolonged cold conditions and shorter growing season in higher elevations likely add a temporal aspect to the commonly used elevational gradient approach, which is not often considered.
4. MBC and enzymatic potential within the rhizosphere were high across all plant species in the alpine meadow during the winter sampling, despite notable differences in microbial community composition. In contrast, winter rhizosphere communities did not differ between plant species in subnival locations, although one species, *Oxyria digyna*, demonstrated much higher microbial activity along with higher NSCs and root N, suggesting some alpine plant species may acquire nutrients through microbial interactions during snow-covered winter periods.
5. *Synthesis*: This study provides the first look at the annual phenology of multiple alpine plant species and their associated rhizosphere microbiome. Our results demonstrate that seasonal microbial dynamics are highly influenced by abiotic factors (soil and microclimatic conditions), but plants are able to modulate these conditions through growth and nutrient acquisition strategies. Taken together, seasonality and independent plant species effects cannot be overlooked when assessing habitat nutrient cycling and ecosystem stability.

## 1. Introduction

Plant-microbial interactions predominate terrestrial ecosystems and drive the ecological stability and functioning of these habitats (Hassani et al., 2018). Within the rhizosphere, the biogeochemical cycling of essential elements, such as carbon (C), nitrogen (N), and phosphorus (P), is dependent upon the facilitation, cooperation, and competition between different trophic levels (Čapek et al., 2018; Lambers et al., 2009). Therefore, the coexistence of microbial communities and multiple plant species must rely on spatiotemporal niche differentiation related to, in part, seasonal changes in nutrient availability and plant-specific phenology (Ke and Wan, 2020). These patterns likely lead to a nuanced microbial phenology, including shifts in soil biological parameters (microbial biomass, extracellular enzymatic activity, respiration, etc.) and microbial community structure (Bonato Asato et al., 2023). Alpine habitats, characterized by a cold climate, prolonged snow cover, short growing seasons, and minimal soil development (Körner, 2021), demonstrate intense seasonal dynamism and temporal nutrient limitation, providing a framework of abiotic factors which may accentuate ecological relationships within and between trophic levels. Furthermore, these ecosystems are especially affected by global climate warming, experiencing accelerated temperature increases (Nigrelli and Chiarle, 2023), shifts in plant distributions (Dolezal et al., 2016; Xu et al., 2023), and an overall ‘greening’ (Wookey et al., 2009). As alpine meadow plants continue to expand upward in elevation into subnival zones (Sklenář et al., 2021; Steinbauer et al., 2018), a concomitant advancement in soil development and microbial communities occurs (Angel et al., 2016; M. Rama Krishna et al., 2020; Ruka et al., 2023). However, a seasonal assessment of plant-microbial interactions across multiple plant species within these changing ecosystems is particularly lacking and would impart insights into the effect of species-specific plant phenology upon microbial physiology and community dynamics.

Within alpine environments, the aboveground growing season is constrained by snowfall, cold temperatures, and drought (Jonas et al., 2008; Möhl et al., 2022) which necessitates the efficient growth and acquisition of nutrients during favorable snow-free conditions (Bilbrough et al., 2000; Körner, 2021; Wang et al., 2020). In contrast, the stable environmental conditions provided by winter snow cover, such as high soil moisture and temperatures above freezing (∼0 °C), have been shown to promote peak microbial biomass measurements in winter and early spring (Brooks and Williams, 1999; Schmidt and Lipson, 2004). Upon snowmelt, a major die-off of these soil microorganisms occurs with the changing environmental conditions, potentially due to osmotic stress and freeze/thaw events (Brooks et al., 1998; Edwards et al., 2006). In conjunction with a large influx of inorganic N made available through melting snowpack, a wide availability of C, N, P and micronutrients derived from plant litter and microbial necromass during spring leads to a pulse of biogeochemical activity, including an increase in microbial extracellular enzymatic activity (Rindt et al., 2023). As the snowpack disappears, N-availability also decreases, resulting in an N-limited environment (Bowman et al., 1993) where plants and soil microbes persist and access nutrients through biotic interactions or phenological processes (Broadbent et al., 2024). During the warm, snow-free months, much of the biogeochemical activity is occurring within the rhizosphere, as plants harbor specific microbial communities and provide a source of carbon through rhizodeposition, which promotes microbial activity (Kawasaki et al., 2021). This dependence on plant-provided photosynthates suggests a close interconnection between aboveground plant phenology and belowground microbial phenology. However, little is known about how closely different plant species or growth strategies are connected to soil microbes, particularly in terms of their interactions with specific microbial groups such as bacteria and fungi.

Amongst alpine plants, temperature and nutrient availability are thought to constrain growth, requiring individual strategies to maximize efficient nutrient utilization and storage (Körner, 2003; Macek et al., 2012). Depending on plant growth forms or functional groups, such as forbs, grasses, or cushions, plant species allocate biomass, and thus nutrients, to specific organs like roots, stems, and leaves for survival and competitive advantages in harsh alpine environments (Doležal et al., 2024). Subsequently, the seasonal distribution of nutrients (C, N, and P) throughout plant tissues is temporally important as plants must perform photosynthesis, biomass production, and storage compound production during a short optimal window of daylength and hospitable temperatures (Liu et al., 2023; Wang et al., 2020). Nitrogen, stored in plant tissues as nitrate, free amino acids, and soluble proteins, is commonly transported from roots to stems and leaves during the growing season for cell synthesis and carbon fixation as an essential component for the ribulose bisphosphate carboxylase/oxygenase (Rubisco) enzyme (Gloser, 2002). While C is used for structural compounds, including cellulose and lignin; plants also utilize and accumulate non-structural carbohydrates (NSCs; starch, fructans, sucrose, etc.) for a myriad of purposes such as osmoprotection, long-term storage during winter, and rhizodeposition (Chlumská et al., 2022, 2014; Kawasaki et al., 2021). However, although the seasonal patterns of NSCs have been observed in selected species (Liang et al., 2019; Signori-Müller et al., 2022), an investigation looking at the seasonal coupling of microbial community dynamics and plant NSC concentrations is currently missing from the scientific literature. Taken together, these plant-specific nutrient allocation strategies represent inconspicuous temporal differentiation occurring between plant species, which likely plays an important role in rhizosphere microbial dynamics.

The phenology of alpine microbial communities is influenced by abiotic and biotic factors, including seasonal changes in climate, edaphic conditions, and plant interactions, yet a comprehensive study remains to be performed to quantify the relative impact of these ecological facets. While the effect of climate and soil on microbes cannot be understated, plant-specific phenology likely interacts with and modulates these abiotic factors, influencing microbial physiology and community composition (Fu et al., 2002; Thoms and Gleixner, 2013) as a means of coexistence within plant communities (Ke and Wan, 2020). Therefore, in this study we sought to disentangle the factors determining seasonal soil biological parameters and microbial community composition across an elevation gradient in the Central Eastern Alps within the rhizosphere of 8 different plant species (Figure 1). The four sampling localities, which spanned a gradient of 600 m of elevation. (2200-2800 m.a.s.l) and encompassed two habitats (alpine meadow and subnival zone), were sampled during the four seasons throughout 2022-2023, including the late snow-covered months. Through climatic dataloggers and soil physicochemical measurements, we captured the seasonal changes in temperature, soil moisture, and soil chemistry along an elevation gradient reminiscent of long-term alpine succession from recently deglaciated terrain (subnival zone; 100-200 years old) to long exposed environments (alpine meadow; >1000 years old). From the rhizosphere of the selected alpine plants, we quantified microbial C (MBC) and N (MBN) content, extracellular enzymatic potential, and performed amplicon sequencing of bacterial and fungal communities to understand the seasonal changes occurring in microbial consortiums at physiological and taxonomic levels. Furthermore, we present a detailed examination of plant nutrient concentrations (C, N, and P) in leaves and roots, NSCs in roots, and leaf isotopic signatures of ^13^C and ^15^N throughout their phenological stages to reveal potential trends and interactions between plant nutrient allocation strategies and soil microbial dynamics. From this experimental design, we aimed to explore three questions: 1) How does microbial phenology differ between these alpine habitats (alpine meadows and subnival zones) in the relative impact of abiotic (climate and edaphic) and biotic (plant species) factors? 2) Which plant-specific traits are associated with seasonal changes in soil biological parameters and microbial community composition? 3) Do plants sustain a rhizosphere effect, in differentiating microbial communities and soil biological parameters, in both habitats during winter?

**Figure 1.**
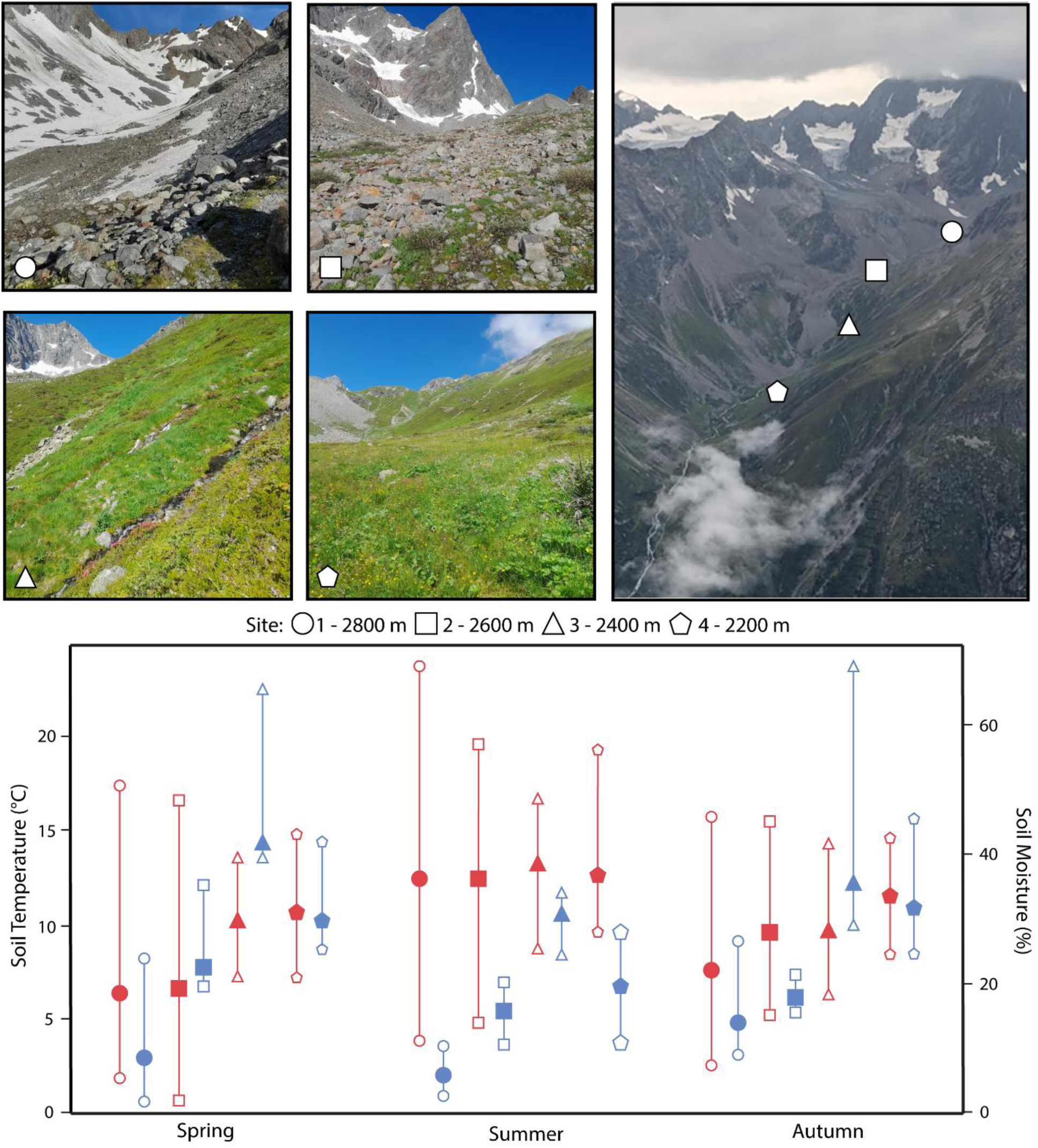
Study sites from the Pitztal region of the Central Eastern Alps spanned an elevation gradient of 2200-2800 m from two habitats, subnival zone (1 & 2) and alpine meadow (3 & 4). Maximum, minimum, and mean values of soil temperature (red) and soil moisture (blue) were recorded at each site using Tomst dataloggers across the growing season.

## 2. Methods

### 2.1 Site description

The study was performed in a valley extending east of Watzespitze (47.00443, 10.79955), a summit in the Tyrol region (Austria) of the Central Eastern Alps near the village of Plangeroß (elevation: 1612 m, mean annual temp: −0.8 °C, annual rainfall: 1320 mm). The growing season is restricted to May-September, depending on snow cover. The warmest and rainiest month is July, averaging 8.9 °C and 158 mm, respectively. Bedrock in the region is dominated by gneiss and mica schists (Schraml et al., 2015). The studied elevation gradient spans from 2200 – 2800 m.a.s.l, starting in an alpine meadow ecotype and ending in the subnival zone of a receding glacier Plangeroßferner (Fig.1). The uppermost locality (Site 1 - 2800 m) is near Kaunergrathütte, where recently deglaciated terrain results in a sparse vegetation community and fine-grained, undeveloped soils. The second highest location occurs within the lower range of the subnival zone (Site 2 - 2600 m), where the terrain becomes smooth, post-glacial hills and pockets of vegetation occurring between the steep, rocky screes on both sides. The second lowest sampling location (Site 3) is at 2400 m, in a steep, vegetated slope with a southern aspect just below the transition from alpine meadow to the subnival zone. The lowest sampling location (Site 4 - 2200 m) is characterized by flat, vegetated meadows, including grasses, forbs, and shrubs, with steep, rocky screes on both sides. In the two lower localities (classified as alpine meadows) snow cover persists typically until late May – early June while in the upper two localities (subnival zone) it may remain until late June – early July. To observe seasonal changes in climatic variables, dataloggers (Tomst™) were continuously recording soil temperature (°C), ambient air temperature, and soil moisture (%). The measurement took place from October 2021 to May 2023 at the four sampling locations.

### 2.2 Field sampling

Sampling efforts occurred during the growing season of 2022 and the following late winter in 2023. At each of the four sites, three replicates of selected dominant plant species (four to five species at each site) were collected in spring, summer, and autumn for roots and rhizosphere soil sampling within a radius of 20 m from the dataloggers. Plants were dug out and excess soil from the roots was shaken off into a sterile bowl for rhizosphere soil samples (∼20 g). Height (cm) of sterile and fertile shoots was measured for each individual plant before washing the plant in a nearby glacial stream and placing them in individual plastic bags. A composite bulk soil sample (100 g) from three non-vegetated patches was also collected at each locality for soil physicochemical analyses. Sub-samples of bulk soil and rhizosphere soil were divided into two samples for chloroform fumigation (∼10 g) and microbial enzymatic potential (∼5 g) analyses. Sampling occurred over two days during five sampling campaigns: Spring (14-15 June 2022), Summer (19-20 July 2022), Autumn (12-13 September 2022), and twice in late Winter (21-22 April 2023 at the lower site 4; 18-19 May at the upper site 1), while plants were still snow covered. The fourth sampling campaign was split into two periods to accommodate the varying snowmelt rates. Due to the physical challenges of winter sampling, only two sites were sampled during the winter season. All samples were kept below 4 C° using dry ice until arriving at the separate institutes (see 2.3, 2.5, and 2.6.1) for subsequent analyses.

### 2.2 Soil physicochemical analyses

Composite bulk soil samples were promptly transported to the Institute of Botany in Trebon, Czech Republic and stored at –20 °C until processing for further analyses. Then, soil samples were oven-dried at 80 °C, until reaching a stable weight, ground in a mortar and sieved to a 2 mm fraction after removing the roots. From the ground and sieved samples, the following measurements were performed. Bulk and rhizosphere soil pH was measured in a 1:5 (soil: deionized water) soil solution. Soil organic matter content (OM; %) was quantified by dry combustion at 450 °C for 5 h. Soil texture (% > 0.5 mm diameter) was measured via wet sieving with higher values indicating coarser soils. Total N (TN; mg/kg) and ammonium content (NH_4_^+^-N; mg/kg) were measured according to Sparks (1996), using a Lachat QC 8500 automated injection analyzer - three-channel FIA system and Thermo Scientific Eager Xperience (Eagersmart 2016) for signal detection. Total P (TP; mg/kg) was quantified following the U.S. EPA Method 365.4 and Standard Methods 4500–P G. The major cations (Ca^2+^, Mg^2+^, K^+^, Na^+^) were quantified by flame atomic absorption spectrometry (ICP-MS) (ICP-QQQ, Agilent Technologies, Tokyo, Japan) (Kopáček et al., 2004). Additionally, to measure the seasonal input of nutrients via soil hydrological processes, Plant Root Simulator (PRS®) (Western Ag Enterprises, Inc., Saskatoon CA) anion and cation probes were inserted 10 cm into the soil at each locality (Qian and Schoenau, 2002). PRS probes contain either a negatively (resin sulfonic acid; SO_3_^-^) or positively (resin qauternary; NH_4_^+^) charged resin that adsorbs cations or anions during the burial period and represents all soil properties (physical, chemical, and biological) which function in nutrient availability. The optimal exposure period for PRS probes to be deployed is 6 weeks, which limited the use to two time periods (Spring-Summer & Summer-Autumn).

### 2.4 Plant selection and specimen processing

From each habitat, four to five herbaceous plants were selected based upon conspicuity and variation in growth forms or functional groups. Within the subnival zone (sites 1 and 2), five plants were selected, including *Doronicum clusii*, *Geum reptans*, *Sibbaldia procumbens*, *Poa laxa*, and *Oxyria digyna*. Four plants were selected from the alpine meadow locations (sites 3 and 4), including *O. digyna*, *Geum montanum*, *Deschampsia caespitosa*, and *Poa alpina*. A subset of plants was collected from the uppermost (site 1: *O. digyna* and *P. laxa*) and lowest sites (site 4: *G, montanum* and *D. caespitosa*) during the winter sampling efforts. Detailed information of the studied plants such as flowering period, preferred habitat, and distribution (summarized from www.actaplantarum.org and www.infoflora.ch) are provided in the supplementary materials (Supplementary Plate A-I).

Plant specimens were processed immediately upon arrival from the field (<24 h after collection). Fine roots from each individual were cut from the primary root stem and stored in 2 ml of LifeGuard™ Soil Preservation Solution (Qiagen). Each individual plant was then separated into leaf, flower, stalk, and belowground parts, dried in an oven at 50°C for 24 h and subsequently milled. Samples were then used for downstream analyses.

#### 2.4.1 Plant tissue nutrient concentrations

The C and N contents, and stable C (δ^13^C) and N (δ^15^N) isotope compositions of plant biomass, were measured to estimate nutrient acquisition and source, water use efficiency, and nutritional status. About 10-30 mg of the milled and homogenized leaves was weighed into tin capsules and placed into a helium-flushed carousel autosampler, then introduced to a Flash Elemental Analyzer (Thermo Scientific) equipped with a single reactor (1020 °C) comprised of chromium oxide for catalytic combustion and copper wire for reduction of NO_x_ to N_2_. A pulse of oxygen was introduced to the reactor simultaneously with the sample for combustion. The sample gases (N_2_ and CO_2_) were transferred via Sulfinert capillaries with helium as the carrier gas through a magnesium perchlorate trap to remove water, and then separated by a PLOT column (Teflon; 80 cm; 1/8-in. OD; Thermo Fisher Scientific part no. 26008243) held at 50 °C. The sample gases were quantified via a thermal conductivity detector (TCD) to determine C and N content (i.e., wt% C and N), and then introduced to a MAT 253 Plus isotope ratio mass spectrometer (IRMS; Thermo Scientific; Bremen, Germany) via the open split of a Conflo IV interface. The raw isotope ratios of the respective gases were determined relative to pulses of laboratory working gases, and the final values are expressed in standard delta notation relative to Vienna Pee Dee Belemnite for ^13/12^C, or Air for ^15/14^N, after correcting for linearity of the ion source, and normalizing to accepted values of laboratory working standards (yerba mate, orchid leaves) and international reference materials (IAEA-600). The precision of the measurement was typically < 0.5‰ for δ^13^C and δ^15^N, both, and < 2 and 0.5% for wt% C and N, respectively. Phosphorus content of plant tissues was measured by colorimetry after perchloric acid digestion (Kopáček and Hejzlar, 1995).

#### 2.4.3 Non-structural carbohydrates (NSC) in belowground organs

##### Ethanol extraction and chromatographic analysis of sugars and sugar alcohols

Milled root samples were used for ethanol extraction and filtration, performed according to Chlumská et al. (2014). All NSC concentrations in this study are expressed as a percentage relative to dry mass. Approximately 100 mg of the dried sample was extracted in 80% ethanol at 83°C for 12 minutes, centrifuged, and the supernatant collected (repeated three times). The remaining pellet was kept for starch enzymatic analysis (see section *Starch analysis*). The supernatant was evaporated and redissolved in distilled water using a shaker. The solution was filtered through nitrocellulose-nitrate membrane filters (Pragopor, retention of particles >0.4 μm). Sugars (glucose, galactose, fructose, sucrose) and sugar alcohols were analyzed using ion-exchange chromatography. Samples were analyzed on Dionex system, using CarboPac PA 10 column with isocratic elution profile using 18 mM NaOH at a flow rate of 1 ml/min at 40 °C, the length of the program was 35 min.

##### Starch analysis

The starch remaining in the pellet after ethanol extraction and centrifugation underwent partial enzymatic hydrolysis and full solubilization using thermostable α-amylase at 100°C. The resulting starch dextrins were further broken down into glucose by amyloglucosidase at 50°C. The glucose concentration was determined spectrophotometrically after glucose oxidase/peroxidase reagent addition. Starch quantities were calculated using the Mega-Calc Excel macro. For additional information, refer to Chlumská et al. (2014) and www.megazyme.com, AOAC Official Method 996.11.

##### Fructan analysis

To extract fructans, approximately 100 mg of the original dried sample powder was boiled for 15 min on a hot-plate magnetic stirrer in 25 ml of distilled water. After cooling, the suspension was filtered through a paper Whatman Grade 1 (retention of particles >11 μm).

A modified version of the method described by Chlumská et al. (2014) was used, incorporating an additional pre-treatment step to remove raffinose-family oligosaccharides (RFOs) and adjusting the temperature for the subsequent enzymatic process. In this pre-treatment, 0.2 ml of the filtered solution was incubated at 40°C for 30 minutes with 50 μl of α-galactosidase (200 U/ml in 50 mM sodium acetate buffer at pH 4.5) to eliminate all RFOs. Next, residual non-fructan NSCs (including poly-, di-, and monosaccharides) were removed by applying sucrase, β-amylase, pullulanase, and maltase. The resulting solution was then incubated at 30°C, and any reducing sugars were reduced using alkaline borohydride and subsequently removed with acetic acid. Other steps for fructan analysis remained the same as in Chlumská et al. (2014). Fructan quantities were calculated using the Mega-Calc Excel macro (www.megazyme.com, AOAC Method 999.03, and AACC Method 32.32.01).

### 2.5 Soil biological parameters

Bulk soil and rhizosphere sub-samples for chloroform fumigation were processed immediately upon returning from the field at the University of South Bohemia in České Budějovice, Czech Republic. Following the protocol described in Vance et al. (1987) (for microbial carbon (MBC) and nitrogen (MBN), soil samples were split into duplicates and half were exposed to chloroform in a desiccator for 24 h. Both sets of samples were extracted via potassium sulfate (4:1 solution – soil) and vacuum-filtered. Extracts were then analyzed for total carbon and nitrogen on a TOC analyzer (TOC/TNM-L, Shimadzu) and values for MBC and MBN were calculated according to Vance et al. (1987) using correction factors of 0.45 and 0.54, respectively.

A second set of sub-samples from bulk soil and rhizosphere samples were stored at −20 °C until enzymatic assays were performed. Soil enzymatic potential was assessed using hydrolase enzymatic assays for five fluorophore-labeled substrates, including β-glucosidase (4-methylumbellyferyl-β-D-glucopyranoside - MUFG), cellobiohydrolase (4-methylumbellyferyl-N-cellobiopyranoside – MUFC), phosphatase (4-methylumbellyferyl-phosphate – MUFP), leucine aminopeptidase (L-leucine-7-amido-4-methylcoumarin – AMCL), and chitinase (4-methylumbellyferyl-N-acetglucosaminide – MUFN). For each sample, 0.5 g was diluted in 50 ml of distilled H_2_O, sonicated for four minutes, and filtered. After testing for the optimal concentration, fluorescence was measured at three time points (30, 90, 150 min after pipetting) using a spectrophotometer (TECAN Infinite F200) with fluorescence intensity filters at 360 and 465 nm for excitation and emission, respectively. Enzymatic potential activity was calculated according to German et al. (2011) and expressed as nmol g^−1^ dry soil h^−1^.

### 2.6 Microbial community composition

#### 2.6.1 DNA extraction and PCR amplification

DNA was extracted using the phenol-chloroform extraction protocol (Angel et al., 2012) at the Institute of Soil Biology and Biogeochemistry in České Budějovice, Czechia. Briefly, for DNA extraction, 0.1 to 0.5 g (wet weight) of root and soil samples were used, depending on the amount available. The full protocol can be found under (Angel et al., 2021). DNA concentration was quantified by Quant-iT™ PicoGreen™ dsDNA Assay Kit (Thermo) and diluted to the range of 1-10 ng μl^−1^ for PCR amplification. Bacterial 16S rRNA gene was amplified using the primers 515F-mod and 806R-mod located in the V4 region (Walters et al., 2015). The ITS2 region for fungal sequences was amplified using the fITS7 (Ihrmark et al., 2012) and ITS4ngsUni primers (Tedersoo and Lindahl, 2016). Primer barcoding was done in a two-step process according to Naqib et al. (2019) based on the Fluidigm Access Array (Fluidigm). For this purpose, the abovementioned primers were included in library preparation synthesized with either the CS1 linker (forward primers) or CS2 linker (reverse primers) on their 5’ ends. For each season, no-template PCR controls (NTC) and blank extractions were included.

#### 2.6.2 Molecular data curation

Library construction and sequencing (253 bp) were performed at the Genome Research Division of the University of Chicago, Illinois, USA in three batches (Spring; Summer and Autumn; and Winter) on an Illumina MiniSeq sequencer (Illumina). For all raw reads, Cutadapt (V3.5) (Martin, 2011) was used to trim the primer and linker regions. For the 16S rRNA gene sequences, quality filtering, merging, denoising, and chimera removal were carried out using the DADA2 pipeline (Callahan et al., 2016) in R v4.4.1 (R Core Team, 2021), with the following non-standard filtering parameters: maxEE = c(2, 2) in the filterAndTrim function and pseudo pooling. Taxonomic assignment was done against the SILVA V138 database (Quast et al., 2013), and sequences not classified as bacteria or archaea were filtered out. Heuristic decontamination was done using the decontam R package (Davis et al., 2018), and unique sequences were compiled to an amplicon sequence variant (ASV) table. A total of 21,894 ASVs (4,981,247 total reads) were included in subsequent analyses after applying a prevalence threshold of >1% and a minimum library size of 2,500 reads (median 22,752) (n = 173). Raw ITS2 reads were inspected using FastQC (v0.12.1) and MultiQC (v1.22) and processed using the PIPITS pipeline (v3.0) using a 97% identity threshold for OTU clustering (Ewels et al., 2016; Gweon et al., 2015).

Subsequently, samples below a minimum library size of 2,000 reads (median 20,650) were removed and OTUs in the remaining dataset (n = 187 samples) were taxonomically classified using BLASTn against the UNITE database (v9.0). Hits with an e-value > 1e-50 and percent coverage < 80% were discarded while the taxonomic assignment of passing OTUs was truncated to genus-, family-, order-, and class-level based on 90%, 85%, 80% and 75% sequence identity thresholds, respectively. After removal of singletons and prevalence filtering using a 1% threshold a total of 5,717 fungal OTUs (4,174,911 reads) were kept for further analysis.

Functional classification of 2,669 fungal OTUs was done based on genus level taxonomy and primary lifestyle in the FungalTraits database (v0.0.3) (Põlme et al., 2020). The processed reads and metadata were ported into the R package Phyloseq (McMurdie and Holmes, 2013) for downstream analysis. Alpha diversity metrics (Shannon-Weiner Index and Simpson Index) of bacterial and fungal communities were calculated from the curated OTU and ASV tables using the ‘diversity()’ function (R package: vegan).

### 2.7 Statistical Analyses

As a preliminary step, a principal component analysis was performed using the seasonal microclimatic and soil physicochemistry data to confirm the habitat classification of the four sampling locations based on clustering (Supplementary Figure 1). To assess the seasonal changes in measured plant traits, soil biological parameters, and microbial alpha diversity, linear mixed effect models (R package: lmer4) were used with the site as a random factor to account for repeated sampling. Log_10_-transformation was utilized according to its efficacy in improving normality and variance tests. Models were first applied to all samples across the four sites with season, plant species and habitat as explanatory variables. Subsequently, the tests were applied separately within habitats to observe differences in seasonal changes between alpine meadow and subnival zone sites. P-values for the linear models were adjusted using false-discovery rate correction. Post-hoc pairwise Wilcoxon rank sum tests were used to determine significant differences between specific seasons and plant species.

To determine the coupling of plant traits with soil biological parameters (MBC, MBN, extracellular enzymes), microbial alpha diversity, and microbial beta-diversity (fungal and bacterial separately), Mantel tests were performed among individual plant species. Prior to the analyses, plant traits, soil biological parameters, and microbial alpha diversity were scaled by root-mean-square transformation while microbial beta-diversity data was normalized by Hellinger standardization. Distance matrices were then calculated with Euclidean distances (R package: vegan) and used for mantel tests. Then, forward selection (package: adespatial) was conducted to assess which plant traits were most closely related to changes in different soil/microbial parameters.

Seasonal changes among fungal and bacterial communities were assessed using Euclidean distance matrices of Hellinger-standardized data for PCoA analyses (“prcomp”, R package: stats). Ordinations were performed within individual plant species across the season and separately for each season among the two habitats. Clusters based upon the 95% confidence interval were generated for seasons or plant species using the “fviz_pca_biplot” function (R package: factoextra). Additionally, the five OTUs contributing most to the ordination were displayed in the ordinations. The ‘adonis’ function was used to conduct permanova analyses (R package: vegan) with 999 permutations for each ordination to determine the effect of season, plant species, and site for the respective analysis.

To develop plant-specific phenology plots of the most correlated plant and soil microbial variables, spearman rank analyses were performed with the forward-selected plant traits as explanatory variables and for each respective group of variables. For bacterial and fungal communities, the five ASVs (prokaryotes) or OTUs (fungi) most contributing to seasonal ordinations were tested for correlations with plant traits. Plant and soil microbial variables found to have a significant relationship for each plant species are listed in the supplementary material (Supplementary Plates A-I), however, only relationships <0.01 were visualized in Figure 6. Means of the selected variables were scaled from 0 to 1 to normalize y-axes for aboveground plant variables and belowground soil/microbial variables.

Last, variation partitioning was performed amongst all plants species within each habitat to quantify the relative effect of climate, edaphic conditions, and plant phenology throughout the growing season (Spring, Summer, Autumn) upon soil biological parameters and microbial communities. Measured climatic variables were extracted from Tomst dataloggers and included maxima, minima, and ranges for soil, surface and air temperatures (°C) and soil moisture from the preceding seven days before sampling events. Edaphic variables included plot-based physicochemistry measurements and individual rhizosphere soil measurements which included extractable C, N, C:N_mol_, and rhizosphere soil pH. To limit the number of explanatory variables from each group, forward selection was performed on root-mean-square-scaled values to identify the two most important variables. Analyses were subsequently performed with the two forward-selected variables from each group (climate, edaphic, plant) for soil biological parameters, microbial alpha diversity, bacterial composition, and fungal composition in each habitat using the “varpart” function (R package: vegan).

## 3. Results

### 3.1 Plot-based environmental conditions

Among edaphic and climatic variables, differences were observed between the subnival and alpine meadow locations. Across the seasons, OM, TN, NH_4_^+^-N, TP, were observed to be higher in the alpine meadow while pH was generally higher in the subnival zone (Table 1.) Other measured edaphic variables, such as texture, PO_4_^3-^, Mg^2+^, Ca^2+^, and K^+^, showed more variability between habitats. Within the Alpine meadow sites, pH (Summer), OM (Spring), TN (Spring), PO_4_^3-^ (Summer), peaked uniformly. Subnival locations also demonstrated maximum values for texture (Summer), NH_4_^+^-N (Spring), PO_4_^3-^ (Summer), Mg^2+^ (Spring) in the respective seasons. However, winter sampling only occurred in sites 1 and 4, limiting our conclusions regarding habitat-level trends in seasonal soil parameters.

**Table 1.** Maximum and minimum values of seasonal soil chemical and physical properties across four sites along an elevation gradient in Pitztal, Austria. All four sites were visited during spring, summer, and autumn while two sites (1, 4) were also visited during the snow-covered winter period (*).

| Habitat | Subnival Zone |  | Alpine Meadow |  |
| --- | --- | --- | --- | --- |
| Site | 1 * | 2 | 3 | 4* |
| Elevation | 2800 | 2600 | 2400 | 2200 |
| pH | 6.65 (Win)<br>5.92 (Aut) | 6.01 (Spr)<br>5.63 (Sum) | 5.12 (Sum)<br>4.93 (Aut) | 5.73 (Sum)<br>5.47 (Aut) |
| OM (%) | 3.0 (Aut)<br>1.1 (Win) | 7.0 (Spr)<br>2.4 (Aut) | 34.7 (Spr)<br>16.8 (Aut) | 35.7 (Spr)<br>4.2 (Aut) |
| Texture<br>(% >0.5 mm) | 62.45 (Sum)<br>39.51 (Win) | 55.26 (Sum)<br>48.20 (Aut) | 60.49 (Sum)<br>41.75 (Aut) | 45.10 (Win)<br>32.67 (Aut) |
| TN (mg/kg) | 494.1 (Aut)<br>262.0 (Spr) | 3,649.5 (Spr)<br>573.0 (Sum) | 8,217.5 (Spr)<br>3,933 (Aut) | 10,322.1 (Spr)<br>1,370 (Aut) |
| NH<sub>4</sub><sup>+</sup>-N<br>(mg/kg) | 1.93 (Spr)<br>1.11 (Win) | 5.83 (Spr)<br>3.45 (Sum) | 14.32 (Aut)<br>9.01 (Spr) | 56.38 (Win)<br>5.61 (Aut) |
| TP (mg/kg) | 787.62 (Win)<br>523.27 (Sum) | 635.94 (Spr)<br>408.01 (Aut) | 1,018.73 (Spr)<br>810.41 (Aut) | 1,138.36 (Win)<br>887.24 (Sum) |
| PO<sub>4</sub><sup>3-</sup> (mg/kg) | 17.49 (Sum)<br>11.83 (Aut) | 14.88 (Sum)<br>8.81 (Aut) | 8.45 (Sum)<br>1.82 (Spr) | 28.27 (Sum)<br>8.41 (Aut) |
| Na<sup>2+</sup> (mg/g) | 0.02 (Spr)<br>0.01 (Win) | 0.02 (Spr)<br>0.02 (Aut) | 0.02 (Sum)<br>0.02 (Aut) | 0.02 (Aut)<br>0.01 (Win) |
| Mg<sup>2+</sup> (mg/g) | 0.04 (Spr)<br>0.03 (Sum) | 0.14 (Spr)<br>0.02 (Sum) | 0.14 (Sum)<br>0.05 (Aut) | 0.29 (Spr)<br>0.07 (Sum) |
| K<sup>+</sup> (mg/g) | 0.08 (Win)<br>0.05 (Sum) | 0.22 (Aut)<br>0.05 (Sum) | 0.22 (Sum)<br>0.17 (Aut) | 0.51 (Win)<br>0.14 (Aut) |
| Ca<sup>2+</sup> (mg/g) | 0.30 (Win)<br>0.18 (Aut) | 1.72 (Spr)<br>0.14 (Sum) | 1.17 (Sum)<br>0.21 (Spr) | 2.76 (Spr)<br>0.39 (Aut) |

Soil temperature observations during the growing season peaked in summer for all sites while minimum temperatures were more variable, with subnival sites experiencing lowest temperatures in spring and alpine meadow sites generally being lowest in autumn (Figure 1). Overall, subnival sites experienced a wider range of temperatures, reflecting more diverse microclimatic conditions, whereas alpine meadow sites are more similar in temperature range. Mean temperatures were similar across all sites (∼12°C) in summer, while spring and autumn followed the mean temperature trends expected across an elevation gradient, with the lowest temperatures occurring at the highest elevations and warmer temperatures in the lower sites. Site 3 was commonly the wettest site (>30% soil moisture) across all seasons, alpine meadow sites were wetter than subnival sites. During summer, the soil moisture range was the smallest across most sites, apart from site 2 which had a smaller range during autumn, being consistently around 20%. These patterns demonstrate the wider range of temperatures and drier conditions in upper, subnival habitats while alpine meadows experience more stable conditions with higher soil moisture.

### 3.2 Mixed-effect models of plant species-based parameters

Amongst the measured plant parameters, with location taken as a random factor, half (7 of 14) significantly changed (p < 0.05) across the seasons and all plant species, namely Leaf N, Leaf C, Leaf δ^15^N, Leaf C:N, Root P, and Root C, and Root Fructose (Figure 2, Supplementary Table 1). Furthermore, plant species significantly differed in 12 of the 14 measured plant parameters (Leaf N, Leaf δ^15^N, Leaf δ^13^C, Leaf C:N, Root P, Root C, Root N, Root C:N, Root Fructan, Root Glucose, Root Sucrose, and Root Fructose). However, because of the wide variation and significant interaction of plant species with season (12 of 14 variables; δ^13^C and Root P were not significant), the difference between the habitats was significant only in two out of 14 variables (Leaf N, δ^15^N). For measured edaphic variables across plant species, extractable organic C and extractable N (2 of 4) demonstrated significant seasonal and plant-specific patterns. Soil extractable C:N and rhizosphere pH were found to be mostly affected by plant species which showed differing values throughout the seasons (Supplementary Figure 2) change seasonally only in the subnival locations. These results, taken together, highlight the greater variation of nutrient allocation and growth strategies amongst plants within the subnival zone in comparison to alpine meadow plants.

**Figure 2.**
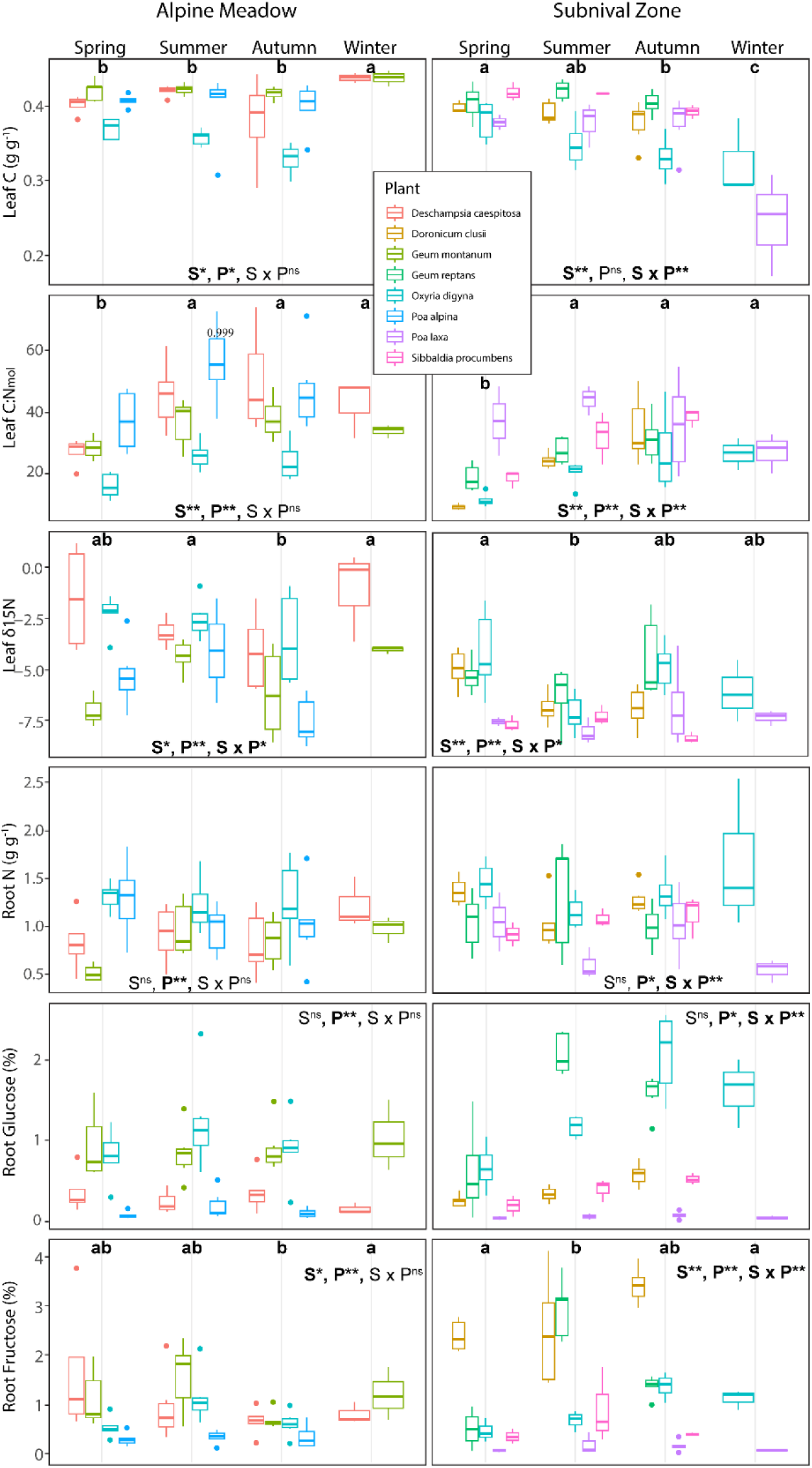
Box plots including linear mixed-effect model results testing the effect of season (S) and plant species (P) individually and their interaction (S x P) upon measured plant parameters within the rhizosphere of 8 different plant species occurring in the alpine meadow and subnival zone. The shown variables were selected based on having the most significant relationships amongst microbial variables when assessing plant species individually. Post-hoc Wilcoxon rank sum tests indicate seasonal differences amongst all plant species.

Almost all soil biological parameters significantly changed across the seasons when combining both habitats (7 of 9, excluding leucinaminopeptidase and microbial C:N; Supplementary Table 1). Of the five enzymatic assays, leucinaminopeptidase and chitinase did not demonstrate a strong plant species effect (Figure 3, Supplementary Table 1). These strong seasonal patterns across habitats appear to be driven by climatic and edaphic conditions present in the alpine meadow locations, as none of the soil biological parameters were found to significantly differ between seasons in the subnival locations (Supplementary Table 1).

**Figure 3.**
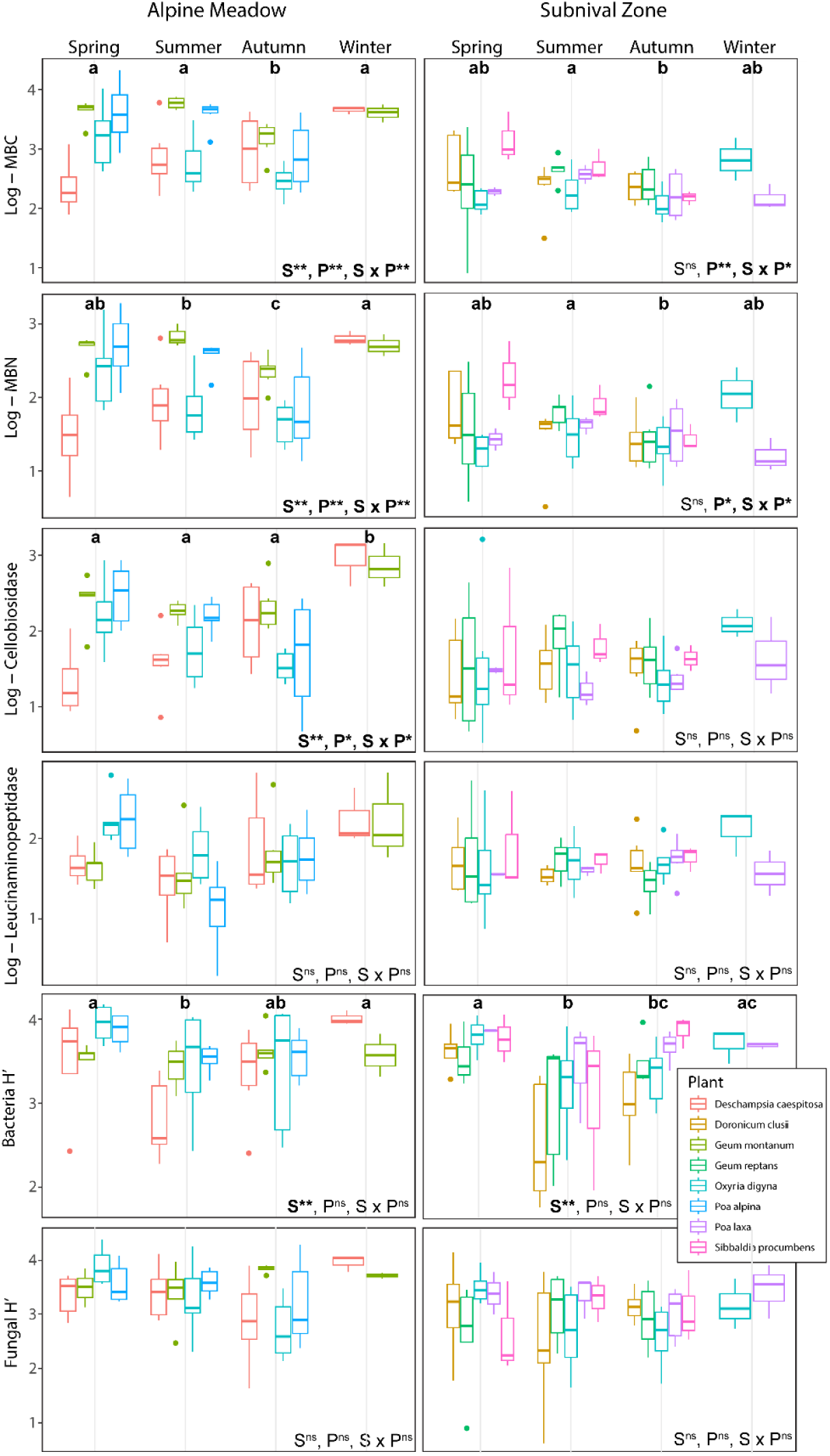
Box plots including linear mixed-effect model results testing the effect of season (S) and plant species (P) individually and their interaction (S x P) upon measured microbial parameters within the rhizosphere of 8 different plant species occurring in the alpine meadow and subnival zone. The shown variables were selected based on having the most significant relationships amongst plant variables when assessing plant species individually. Post-hoc Wilcoxon rank sum tests indicate seasonal differences amongst all plant species. Abbreviations: MBC – microbial biomass carbon, MBN – microbial biomass nitrogen.

However, MBC and MBN differed within the rhizospheres of plant species within the subnival habitat. Across microbial alpha diversity metrices, bacterial ASV richness and Shannon-Weiner (H’) showed a strong seasonal dynamic while fungal OTU richness was primarily associated with plant species and higher richness in the alpine meadow locations (Supplementary Table 1). Apart from fungal OUT richness, not many differences were observed among alpha diversity metrices between habitats, with both habitats primarily showing a seasonal pattern in bacterial communities. Therefore, although plants in higher elevations exhibit stronger seasonal patterns in nutrient allocation and edaphic rhizosphere conditions, these effects are not observed in soil biological parameters nor alpha diversity metrices.

### 3.3 Seasonal patterns in rhizosphere bacterial and fungal beta diversity

Overall, there was more total variation among fungal communities than bacterial communities within the rhizospheres of 8 alpine plant species (Tables 2 and 3). Six of the eight plant species revealed changing fungal communities across the seasons and only one species (*Geum montanum*) did not show a significant effect of location upon fungi (*Sibbaldia procumbens* was only collected at one location). Based on the top five fungal OTUs determining PCoA ordination axes, several fungal species showed similar seasonal associations between plant species. *Microbotryum scozonerae* (Basidiomycota), a plant pathogen, showed a stronger seasonal pattern in *Doronicum clusii* and *Oxyria digyna*, typically becoming more abundant in summer. *Herpotrichia pinetorum*, a common conifer pathogen with root endophytic capability in boreal, mountainous regions, was associated with two plant species, *Geum reptans* and *Oxyria digyna*, in spring and summer (Table 2). An unidentified species within the genus *Lachnum* was associated with *Sibbaldia procumbens* and *Poa laxa* in summer in subnival locations. Last, an unidentified species of genus *Chalara* was found to be more abundant in either summer or autumn of three species (*P. laxa*, *G. montanum*, and *P. alpina).* Permanovas across all rhizosphere and bulk soil samples in separate seasons and habitats revealed that fungal communities were differentiated by plant species in spring and summer within alpine meadow and subnival location, respectively (Figures 4 & 5; Supplementary Table 2). Furthermore, in winter, plant species maintained a significant association with fungal communities in alpine meadow locations, while fungal assemblages became more homogenous during winter within subnival locations. These results confirm that fungal rhizosphere communities in alpine environments are commonly changing with seasons and specific associations exist between multiple plants and fungal species in certain times of the year.

**Figure 4.**
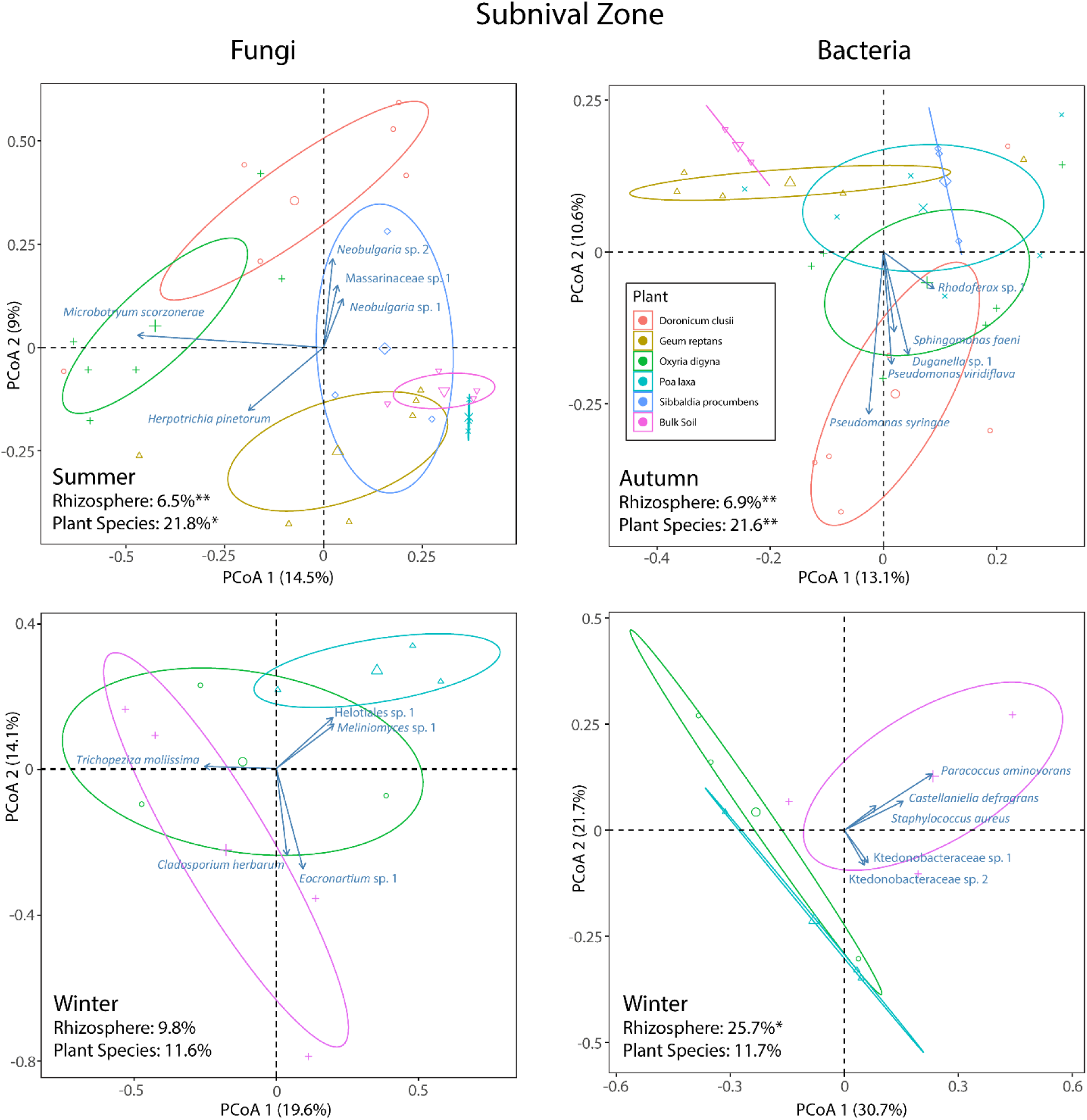
Principal coordinate axes of Hellinger-standardized fungal and bacterial communities of the bulk soil and rhizosphere within the subnival locations; clusters with centroids are indicated by 95% confidence interval. Significance of rhizosphere and plant species compared to bulk soil communities were analyzed using permanova tests with location as a covariate (*p < 0.05, ** <0.01). Summer and autumn are shown for the fungal and bacterial communities, respectively, representing the seasons with the strongest plant species effect upon the rhizosphere composition. Winter is shown for fungal and bacterial communities.

**Figure 5.**
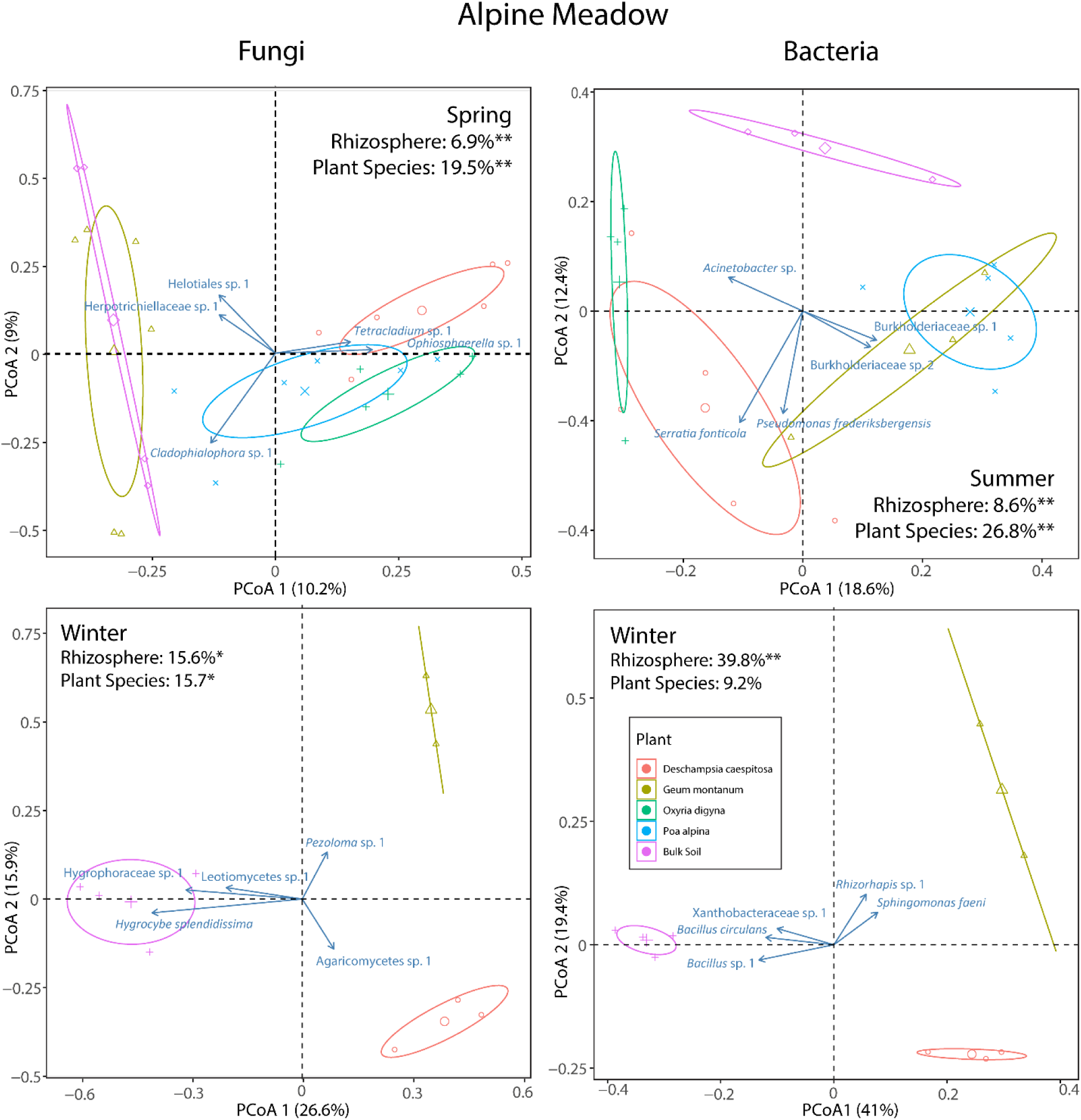
Principal coordinate axes of Hellinger-standardized fungal and bacterial communities of the bulk soil and rhizosphere within the alpine meadow; clusters with centroids are indicated by 95% confidence interval. Significance of rhizosphere and plant species compared to bulk soil communities were analyzed using permanova tests with location as a covariate (*p < 0.05, ** <0.01). Spring and summer are shown for the fungal and bacterial communities, respectively, representing the seasons with the strongest plant species effect upon the rhizosphere composition. Winter is shown for fungal and bacterial communities.

**Figure 6.**
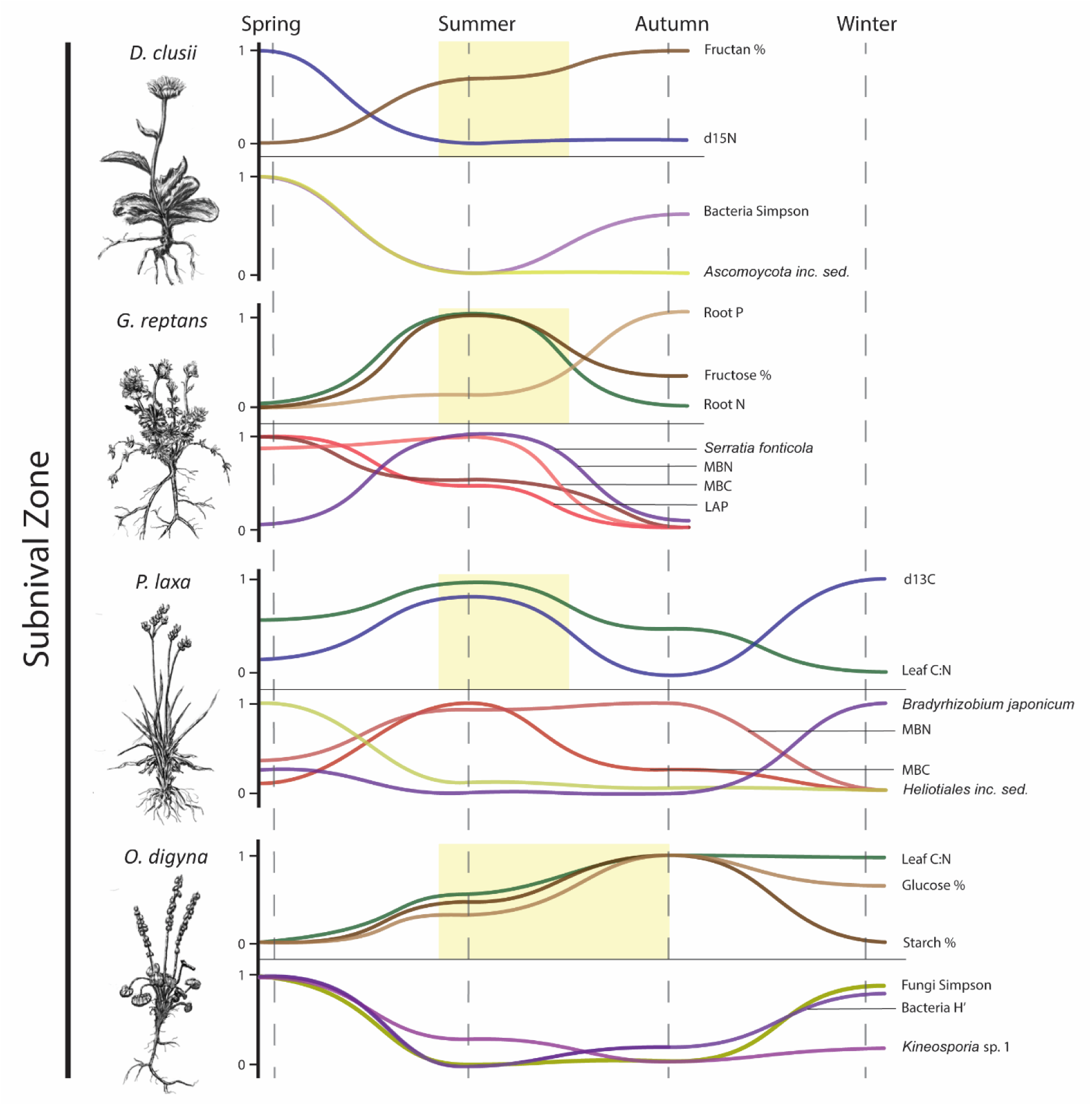
Subnival zone spaghetti plots depicting phenological relationships between scaled means of plant parameters (top) and rhizosphere microbial parameters (bottom) of selected plants. Plant parameters were chosen by forward-selection and further assessed by iterative spearman rank correlation tests to indicate coupling with specific microbial parameters (Supplementary Plates A-I). Plant parameters include leaf nutrients (green), leaf isotopes (blue), and root nutrients (brown). Rhizophere parameters include bacterial diversity (purple), fungal diversity (yellow), and soil biological parameters (red). Typical flowering periods of each plant are visualized by the yellow boxes.

**Table 2.**
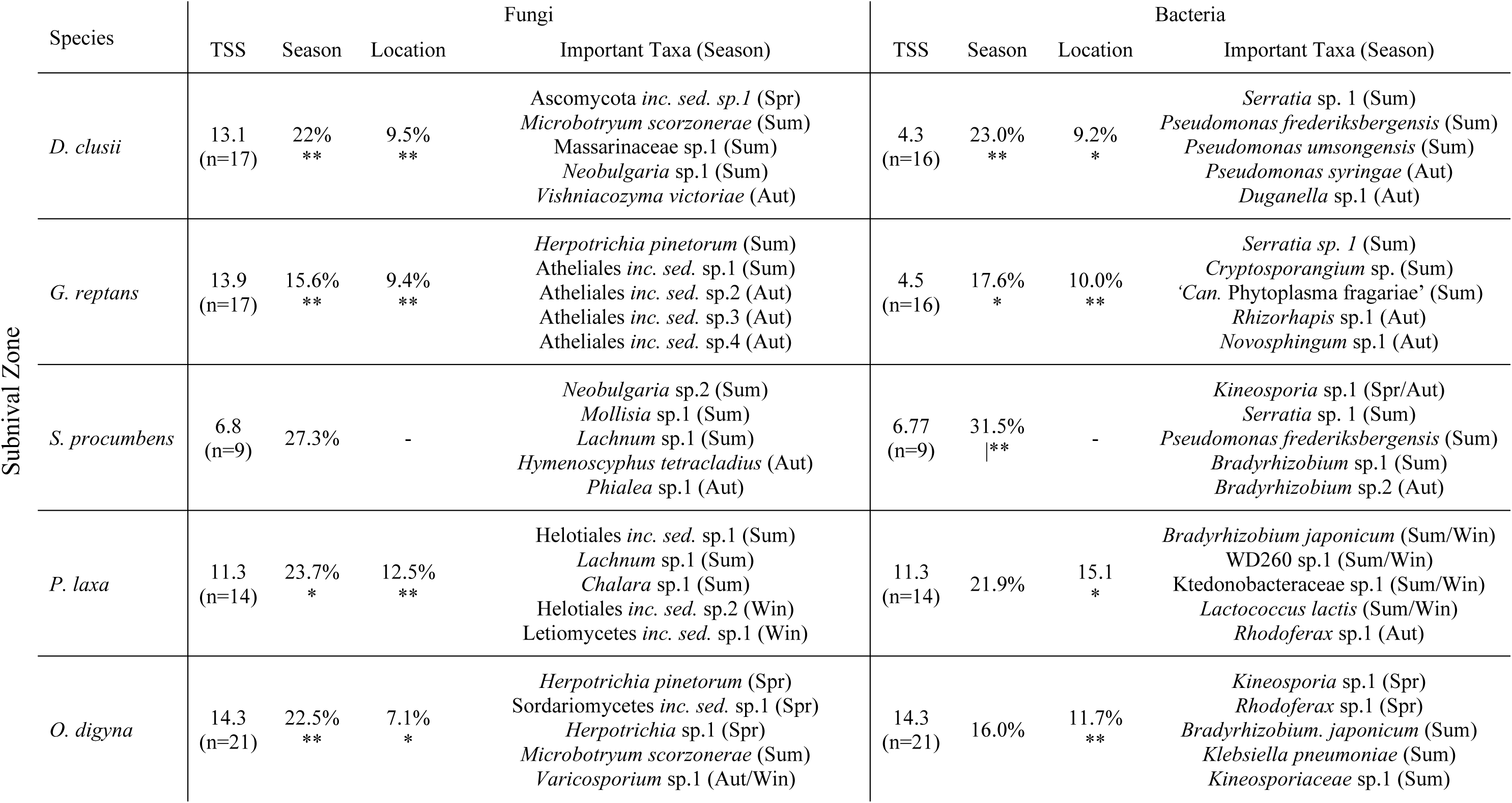
Permanova results of Hellinger-standardized bacterial ASV and fungal OTU composition within rhizosphere samples of several plant species with season and location as explanatory variables within the subnival sites. The proportion (%) of explained variation and significance of each variable is indicated (* p <0.05, ** <0.01) as well as total variation (TSS). Important taxa were determined by identifying the top five species most contributing to seasonal PCA axes.

**Table 3.**
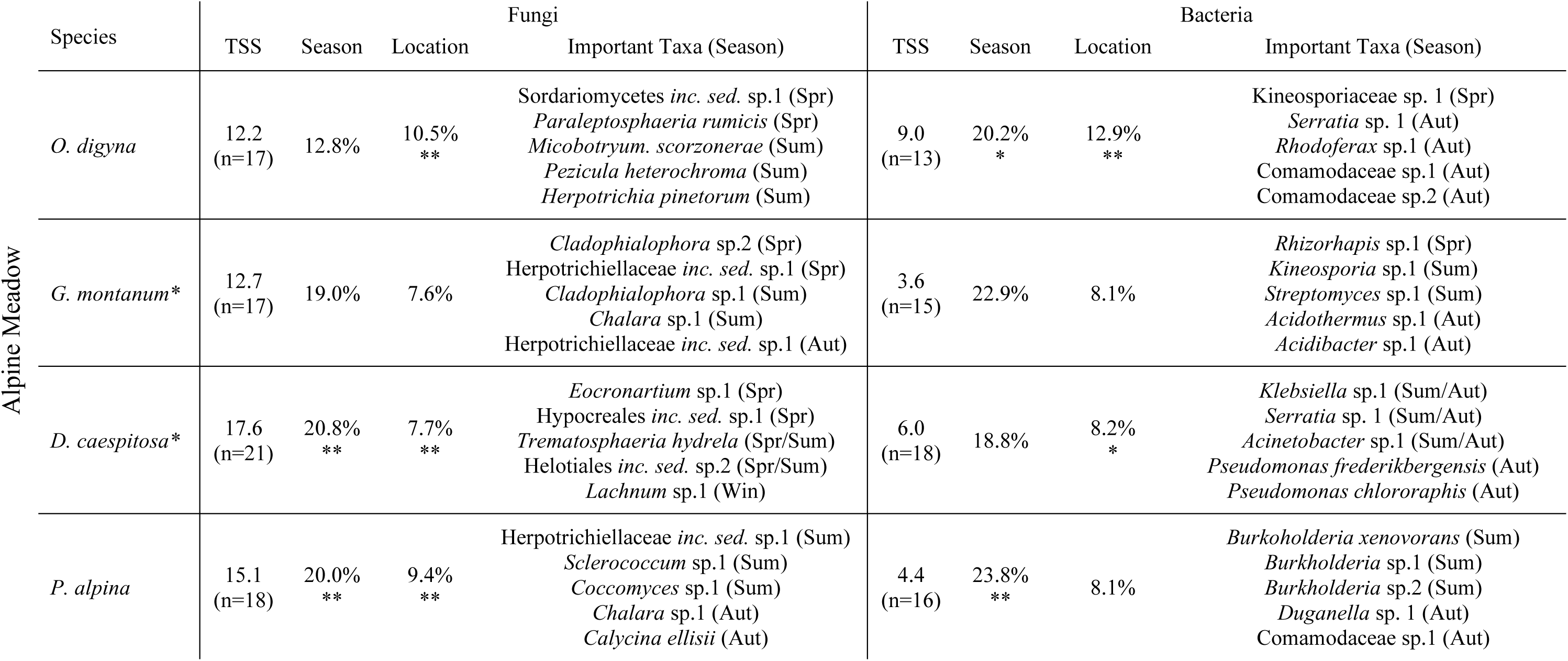
Permanova results of Hellinger-standardized bacterial ASV and fungal OTU composition with season and location as explanatory variables across different plant species within the alpine meadow sites. The proportion (%) of explained variation and significance of each variable is indicated (* p <0.05, ** <0.01) as well as total variation (TSS). Important taxa were determined by identifying the top five species most contributing to seasonal PCA axes.

Amongst bacterial communities, five of the eight plant species had significantly different assemblages across the seasons, while two of the eight species, *Geum montanum* and *Poa alpina,* showed no significant effect of location, as determined by permanova (Tables 2 and 3). *Serratia* sp. 1 was observed in the rhizosphere of five plants across both habitats, becoming more prevalent in summer. *Pseudomonas frederiksbergensis* also demonstrated seasonal preferences in three plant species (*D. clusii*, *S. procumbens*, and *D. caespitosa*) in either summer or autumn. Last, *Bradyrhizobium japonicum* was more abundant in summer amongst rhizosphere communities of *P. laxa* and *O. digyna*. In separate permanova analyses across all rhizosphere and bulk soil samples, the greatest differentiation of bacterial communities by plant species occurred in summer at alpine meadow locations and in autumn at subnival locations (Figures 4 & 5; Supplementary Table 2). Additionally, there was a significant difference between bulk soil and rhizosphere bacterial communities during winter in both habitats, yet no significant difference was found between the assemblages between plant rhizospheres. In total, bacterial communities appear to be less associated to specific plants, but perhaps more constrained by the differences between bulk soil and rhizosphere conditions.

### 3.4 Plant parameters associated with rhizosphere microbial variables

Soil biological parameters and microbial diversity were primarily associated with plant parameters specific to each species. Among soil biological parameters, only two species (*Sibbaldia procumbens* and *Deschampsia caespitosa*) were found to have significant correlations with seasonal plant measurements according to mantel tests (Supplementary Table 3). Based on forward-selection, Leaf C, Leaf C:N, and Root P were the three most common plant traits to be associated with soil biological parameters across all plant species. When assessing correlations between microbial alpha diversity and plant parameters, only *Oxyria digyna* displayed a significant positive coupling. For fungal and bacterial beta-diversity separately, six of eight plant species demonstrated positive correlations with plant parameters. Root N, Root C:N and Root Sucrose were the most common plant variables associated with fungal beta-diversity, whereas δ^15^N, Leaf N, and Leaf C were most commonly associated variables with bacterial beta-diversity across plant species (Supplementary Table 3).

Individual phenologies of plant species and their associated rhizophere parameters constructed from significant Spearman rank correlation tests (p < 0.01) demonstrated that the plant parameters which coupled with microbial parameters varied according to plant species (Figures 6 & 7). Amongst plant parameters, Leaf C was the most common nutrient metric to be correlated with soil biological parameters and microbial communities (3 plants), followed by δ^15^N, Root Fructose, Root N, Leaf C:N, and Root Glucose (2 plants). NSCs were found to be commonly coupled with rhizosphere variables, showing significant relationships in five different plant species (Figures 6 & 7). Soil biological parameters and microbial diversity were more uniform, with MBN being the most correlated with aboveground variables (4 plants), followed by MBC (3), leucinaminopeptidase (2), and fungal Simpson index (2). Due to the autocorrelation of MBC and MBN (Mic C:N consistently around 8-10), there was a common relationship between Leaf C and these variables in graminoids and *Geum reptans* (Supplementary Plates). In seven of eight species, specific microbial taxa displayed significant associations with plant parameters (Figures 6 & 7). These findings highlight the importance of plant phenology upon soil biological parameters and microbial communities and the role of NSCs in modulating the transfer of nutrients between trophic levels.

**Figure 7.**
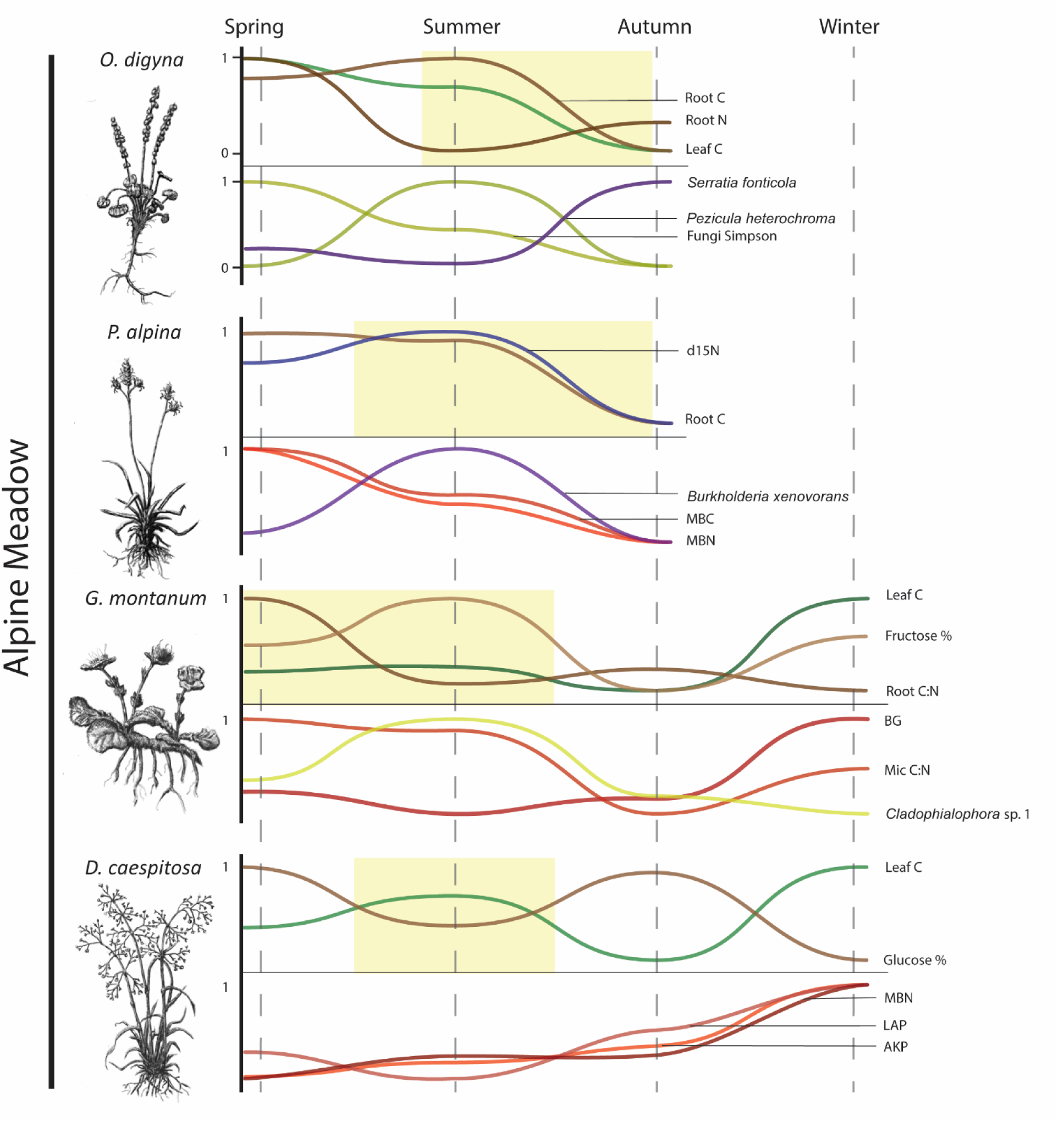
Alpine meadow spaghetti plots depicting phenological relationships between scaled means of plant parameters (top) and rhizosphere microbial parameters (bottom) of selected plants. Plant parameters were chosen by forward-selection and further assessed by iterative spearman rank correlation tests to indicate coupling with specific microbial parameters (Supplementary Plates A-I). Plant parameters include leaf nutrients (green), leaf isotopes (blue), and root nutients (brown). Rhizophere parameters include bacterial diversity (purple), fungal diversity (yellow), and soil biological parameters (red). Typical flowering periods of each plant are visualized by the yellow boxes.

### 3.5 Variation partitioning of abiotic and biotic factors upon soil microbial parameters

The relative importance of climatic, edaphic, and biotic factors showed similarities and differences between habitats and levels of rhizosphere microbial parameters. In both habitats, soil biological parameters (MBC, MBN, and extracellular enzymatic activities) represented the most explained variation (>44%), with soil chemistry (i.e. mainly C and N contents) being the predominant factor (Figures 8 & 9). Microbial alpha diversity demonstrated the second most explained variation (>20%) in both habitats, with microclimatic conditions being the primary factor in the alpine meadow locations while plant parameters were the primary factor in the subnival locations. Among both habitats, total explained variation was higher for bacterial beta-diversity than fungal beta-diversity. Additionally, the explained variation by interactions between the groups of variables were commonly higher than the independent effects of groups. Forward selection analyses within the subnival zone showed a strong effect of ammonium, soil moisture, root N, and some NSCs, such as root fructose and sucrose, upon most of the levels of rhizosphere microbial parameters. In contrast, alpine meadow locations were primarily associated with rhizosphere pH, soil moisture, leaf C, and root starch. Taken together, the forward selection and subsequent variation partitioning analyses show the higher importance of environmental conditions, particularly upon rhizosphere community structure, in subnival locations, while plants likely play a larger role in alpine meadow locations through seasonal microedaphic effects (such as rhizosphere pH).

**Figure 8.**
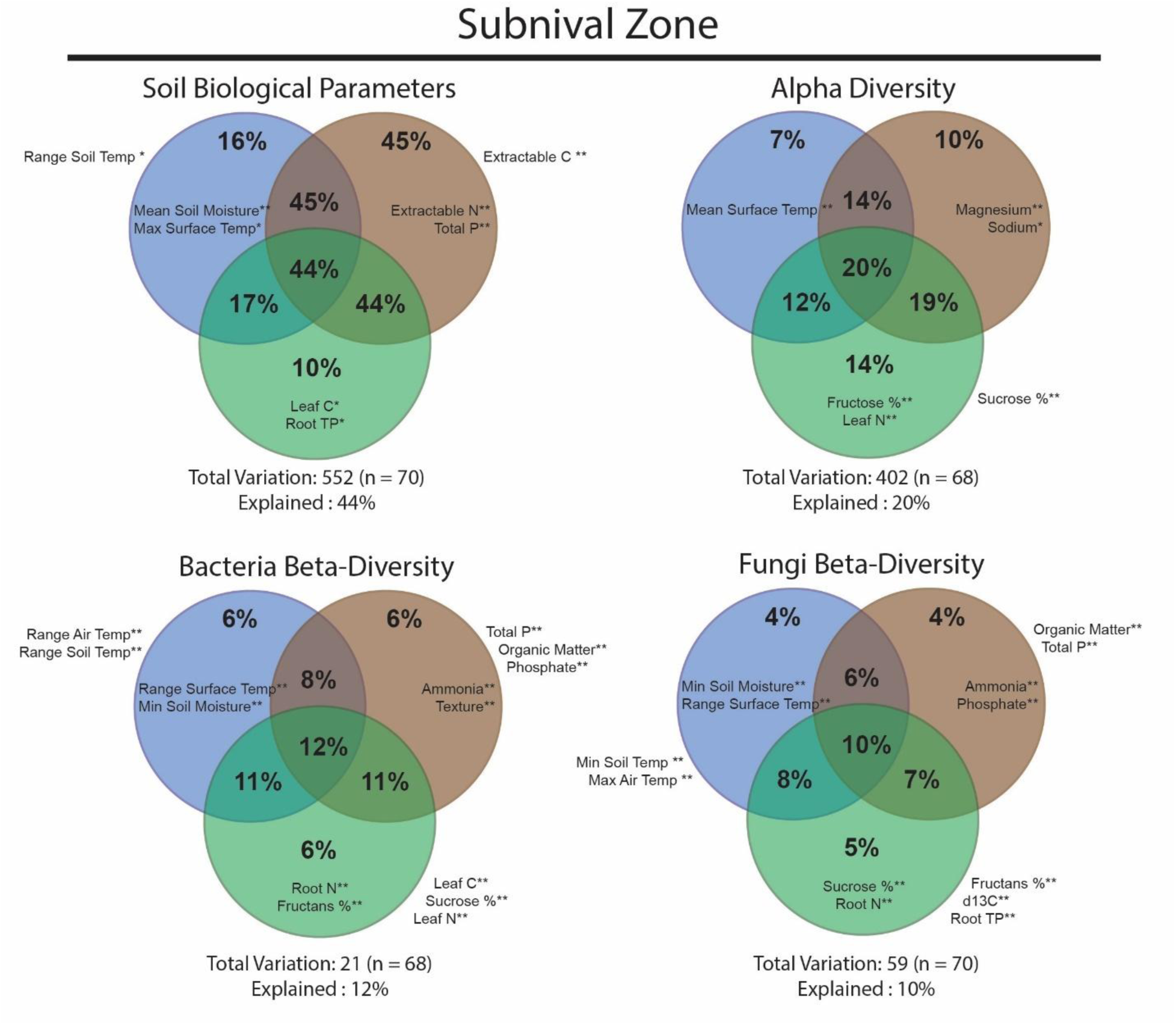
Variation partitioning analyses of rhizosphere parameters measured during the growing season of selected subnival zone plants. Microclimatic data (blue), soil measurements (brown), and plant nutrient concentrations (green) were used as groups of explanatory variables. Up to two variables were included in the analysis (within circle) for each group, based on forward-selection (* p <0.05, ** <0.01).

## 4. Discussion

Understanding the phenology of microbial communities in alpine habitats and their associated biogeochemical processes is essential for predicting the shift in how these ecosystems will function under changing climate scenarios (Broadbent et al., 2024). With strong seasonal dynamics and prolonged snow cover, alpine ecosystems offer an ideal system to study the dynamic interplay of climatic variation, soil chemistry, and plant species upon microbial phenology (Schmidt and Lipson, 2004). This study provides further clarity to the impact of these interactions by assessing the seasonal changes in soil biological parameters (MBC, MBN, and enzymatic potential) and microbial communities within the rhizosphere of eight species in the Central Eastern Alps across two different habitats. By combining plant nutrient measurements with climatic variation and edaphic conditions, rhizosphere microbial parameters were shown to be primarily influenced by plants and their subsequent effect upon rhizosphere pH in alpine meadows (Figure 9); while inorganic N availability (NH_4_^+^-N) and plant N demand played an important role in subnival locations (Figure 8). NSCs were consistently associated with microbial beta-diversity and soil biological parameters, suggesting a potential passive or active carbon storage in plants which may promote or deter microbial activity (Figures 6 & 7). Furthermore, sampling efforts in the late-winter, snow-covered period imply that plant species have differential effects upon biogeochemical cycling and microbial communities even when dormant and non-photosynthesizing (Figures 4 & 5). While these findings are correlational, our comprehensive approach across multiple species, seasons, and habitats reveals microbial phenology is affected by the interaction of abiotic and biotic factors, including collaboration and competition for nutrients in a temporally constrained environment (Jaeger III et al., 1999).

**Figure 9.**
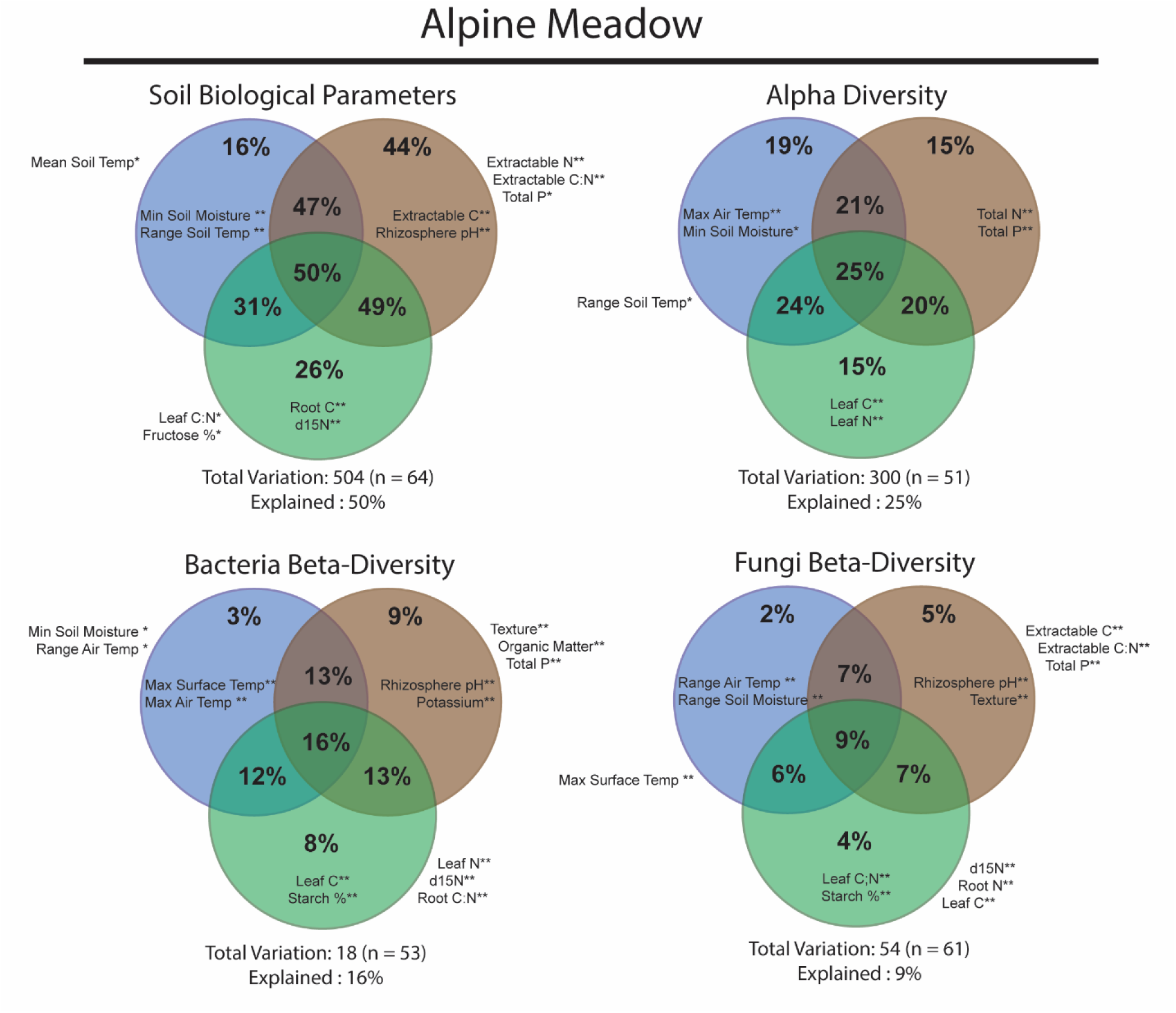
Variation partitioning analyses of rhizosphere parameters measured during the growing season of selected alpine meadow plants. Microclimatic data (blue), soil measurements (brown), and plant nutrient concentrations (green) were used as groups of explanatory variables. Up to two variables were included in the analysis (within circle) for each group, based on forward-selection (* p <0.05, ** <0.01).

### How does microbial phenology differ between alpine meadows and subnival zones in the relative impact of abiotic (climate and edaphic) and biotic (plant species) factors?

Alpine ecosystems undergo strong environmental changes seasonally, from deep snow-covered winters to dry hot summers, leading to predictable pulses in nutrients, such as an influx of N and P during spring snowmelt (Hiltbrunner et al., 2005; Weintraub, 2011). While our study confirms that these seasonal changes in soil chemistry led to predictable increases in cellobiosidase chitinase activity, as well as bacterial alpha diversity (Figure 3, Supplemental Figure 2), in spring and winter, our results also suggest a clear distinction between plant growth strategies and the response of soil biological parameters during the growing season. Within the alpine meadow localities, *Deschampsia caespitosa*, a fast-growing and highly seasonal graminoid (Davy and Taylor, 1975), exhibits pronounced dynamics in such parameters as Leaf C:N, Leaf δ^15^N, and NSCs, which is mirrored by similar strong dynamics in MBC, MBN, and enzymatic potential (Figures 3 & 4). On the other hand, *Geum montanum*, a clonal herbaceous species that is rather aseasonal in its nutrient allocation (with evergreen leaves), shows less seasonality in microbial parameters with consistently high MBC, MBN and enzymatic activity (Figures 3 & 4) throughout the year. Furthermore, the phenology of bacterial and fungal communities differed between these two species. While *D. caespitosa* presented high seasonality in rhizosphere microbial communities, *G. montanum* hosted a more static rhizosphere environment across alpha diversity metrices and fungal and bacterial assemblages (Figure 3, Table 3). Even though these two species often grow intermingled in the studied alpine meadows, their specific growth strategies such as rapid nitrogen acquisition in *D. caespitosa* (Jaeger III et al., 1999) and a steady-state transfer of nutrients through clonal rhizomes in *G. montanum* (Jónsdóttir et al., 1996) show the close, spatial coexistence of plant species may be related to temporal niche differentiation in rhizosphere interactions.

Interestingly, the microbial phenology in the subnival zone is much more subdued than alpine meadows, despite our observations that plants growing there tend to be more seasonal, particularly in δ^13^C and NSCs (Supplementary Table 1). Although individual differences between plant species can be seen in MBC and MBN throughout the season, as well as a decrease in bacterial H’ during summer amongst most rhizosphere communities, the diminished seasonality of soil biological parameters attests to the general lack of organic substrate available (Merino et al., 2016) and the stressful environmental conditions which leads plants to actively store NSCs as a means for cryoprotection and drought stress (Chlumská et al., 2022).

Additionally, the forward-selected variables for variation partitioning consistently included parameters related to N and P (Figure 8), such as ammonium, phosphate, Total P, Root N, and Root P, suggesting nutrient limitation and thus competition between plants and microbes is more prevalent in the subnival zone (Čapek et al., 2018; Jaeger III et al., 1999; Schmidt et al., 2014). Even so, due to our correlative approach, we can only speculate at best upon these interactions, despite the recent progress in understanding the seasonal dynamics of N dynamics in alpine ecosystems (see Broadbent et al., 2024). A key difference between alpine meadow and subnival localities is the phenological delay that occurs at higher elevations. In the Qinghai-Tibetan plateau, Wang et al. (Wang et al., 2014) demonstrated that plant phenology differed in start dates and duration depending on plant functional groups and flowering periods. Within our study, we extend this idea of phenological delay to rhizosphere microbial assemblages by showing that the strongest differentiation of fungal and bacterial communities occurs typically later in the growing season in the subnival zone compared to alpine meadow localities (Figures 4 & 5). The reasons for this are likely many folds. The distinct fungal communities observed in the alpine meadows early in the season may be attributed to the greater temporal separation of flowering periods among plant species (Figure 7), as demonstrated by Liu and Howell (2021). The accumulation of NSCs like starch and fructan during summer and autumn months (Supplementary Figure 3) can shift microbial communities towards oligotrophic species, as suggested by Kawasaki et al. (2021) or plant-specific strategies regarding plant marcescence (Chondol et al., 2024) acting as a stable nutrient pool during senescence for plants and bacterial communities in autumn (Schmidt et al., 2007). However, this temporal divergence in maximum differentiation between bacterial and fungal communities and alpine habitats questions the standard elevational gradient approach, which typically consists of sampling at a single time-period during optimal plant biomass accumulation or flowering (Praeg et al., 2019; Yang et al., 2014; Yao et al., 2017). While it is not reasonable to expect studies to always employ a seasonal approach, it is important for researchers to take into consideration the role of a potential phenological delay amongst multiple trophic levels when using an elevation gradient approach.

### Which plant nutrient parameters are commonly associated with seasonal changes in soil biological parameters and microbial community composition?

At the habitat level, the plant parameters which coupled with changes in belowground phenology differed broadly, with alpine meadow microbial communities being related to plant C dynamics, such as starch, and root or leaf C content, while subnival communities more closely associated with plant N concentrations and fructans or sucrose content (Figure 8, 9). Importantly, these C dynamics in the alpine meadow may potentially lead to changes in the microedaphic conditions in the rhizosphere, such as pH, and effectively influence microbial composition and soil biological parameters (Jones et al., 2009). Nevertheless, the production of insoluble starch granules by *Oxyria digyna* and *Geum montanum* (forbs) in summer and autumn represents a shift towards carbon storage, in contrast to *Poa alpina* and *Deschampsia caespitosa* (graminoids) that produce fructans as storage compounds (Supplemental Figure 3). Yet, even with these similar growth strategies, their microbial communities differed widely and the specific taxa which closely associated with increased root C in summer varied (Figure 7). Admittedly, it is known that sugars are not the primary compounds that influence the rhizosphere but act as a general attractant (Badri et al., 2013), as perhaps seen within our study with *O. digyna*, *G. montanum*, and *P. alpina* demonstrating a positive relationship between root C or fructose content and MBC, microbial C:N, and specific fungal taxa (such as *Pezicula heterochroma* in *O. digyna*) (Figure 7, Supplementary Plates F-I). While not measured in this study, phenolic compounds, known to modulate soil microbiomes (Badri et al., 2013), could explain the stable microbial composition and belowground dynamics of G. montanum, a species known for producing a glycoside, gein, in its’ leaves and roots (Schmidt et al., 2014).

Similarly, within subnival localities, NSCs such as fructans and sucrose played an important role in microbial phenology. Similar to starch, fructans are viewed as long-term storage carbohydrates with additional cryoprotectant effects (Valluru and Van den Ende, 2008), yet their abundance within roots does not show a specifically seasonal pattern, apart from *D. clusii*. Sucrose, on the other hand, a disaccharide product at the end of photosynthesis which is capable of being cleaved into fructose and forms of glucose, is representative of an active photosynthate that is highly sought after by rhizosphere microbes (Chen et al., 2023; Stein and Granot, 2019). While high root sugar excretion has been shown to increase the alpha diversity of microbial communities in engineered *Arabidopsis* plants (Song et al., 2022), the rhizosphere communities within our study showed varying responses to increased simple sugar content, such as *Oxyria digyna* displaying a negative relationship between microbial alpha diversity and glucose content, *Geum reptans* showing a positive correlation with fructose content and *Serratia fonticola* (a known plant growth promoting bacteria) (Kulkova et al., 2024), or *Doronicum clusii* presenting a positive correlation between fructose content and *Pseudomonas syringae* (a known pathogenic bacteria) (Xin et al., 2018). These differing responses in belowground phenology to increased simple sugar content across multiple species demonstrate the contrasting responses of microbial communities when compared to a single model species such as *Arabidopsis*.

### Do plants maintain a rhizosphere effect in both habitats during winter?

The individual effects of plant species upon over-wintering rhizosphere microbial communities have rarely been studied, typically addressing snowmelt dynamics (Rindt et al., 2023) or the effect of snow-removal (Broadbent et al., 2024; Cai et al., 2024). However, the four selected species sampled during the late winter period reveal the potentially homogenizing effect of snow cover, such as the similar values in MBC, MBN, and enzymatic activity in alpine meadows between *Deschampsia caespitosa* and *Geum montanum*, while the subnival species (*Oxyria digyna* and *Poa laxa*) were markedly different in the same belowground parameters (Figure 3). The high amounts of senescent vegetation and soil organic matter content in the alpine meadow likely allows for substantial growth by microbial communities as they consume carbon polymers such as cellulose and chitin (Figure 3) (Schmidt et al., 2007). Despite these similarities in soil biological parameters, the fungal and bacterial communities of *D. caespitosa* and *G. montanum* still differentiated as in other seasons (Figure 5). Interestingly, upon snow removal of the selected plots, *G. montanum* was flush with green leaves, appearing as if during mid-growing season and adding to the concept that some aseasonality exists among alpine plants which is conferred to the belowground community. Contrastingly, the soil biological parameters measured within the subnival localities were notably different between the plants, with *O. digyna* being higher in most measured variables than *P. laxa* (Figure 3). Meanwhile, no significant difference was found between the rhizosphere communities, which suggests that *O. digyna* is promoting the winter microbial communities through simple sugars (e.g. fructose and glucose) during the winter, whereas *P. laxa* stores minimal carbohydrates during the winter. However, at this same late winter period, *O. digyna* is also having increased root N measurements, extending the idea that N uptake can take place during snow-covered periods in alpine environments (Onipchenko et al., n.d.), as in subarctic environments (Larsen et al., 2012). In conjunction with increased microbial activity, nutrient acquisition during a seemingly dormant period would be advantageous for plants to limit competition with microbes in the nitrogen-limited growing season. Although these speculations are not conclusive, they offer testable hypotheses for future multi-species studies to fully understand the mechanisms and prevalence of positive plant-microbial interactions in the winter season.

## Conclusion

This study portrays the associations of plant phenology with rhizosphere microbial phenology of 8 alpine plant species through soil biological parameters and microbial community composition. The distinct seasonal patterns as a result of plant-specific growth strategies, such as rapid nutrient acquisition and growth in contrast with aseasonal clonal growth, demonstrates that microbial communities and biogeochemical cycling can be, in part, explained by seasonal plant nutrient allocation. Seasonal pulses of nutrients, such as during snowmelt and autumn senescence, create key moments for N and P uptake for plants and microbes, yet in the subnival zone, limiting nutrients are potentially tightly linked with soil microbial phenology. One subnival plant, *Oxyria digyna,* perhaps presents a strategy for mitigating this seasonal competition by maintaining high NSCs throughout winter and promoting microbial activity, resulting in higher soil nutrient availability and acquiring N during winter months. However, our study also shows that NSCs have variable impacts upon seasonal microbial parameters, from the accumulation of long-term storage carbohydrates potentially inhibiting microbial activity to simple sugars being associated with decreased microbial alpha diversity or increasing plant growth-promoting bacteria and pathogens. Broadly, we observed a phenological delay between alpine meadows and subnival zones in the peak differentiation of rhizosphere communities among fungi and bacteria. Conclusively, the seasonality of alpine microbial communities and their role in biogeochemical cycling is undoubtedly shaped by their dynamic abiotic environment, yet, as plant species move higher in elevation, their relative and distinct impact upon microbial phenology must be considered.

## Acknowledgements

ATR, JS, RA and KR were supported by the Czech Science Foundation (GA ČR), grant numbers 21-04987S, 24-11954S and RVO 67985939, 60077344. The funders had no role in study design, data collection and interpretation, or the decision to submit the work for publication. We would like to thank Michael und Julia Dobler who manage the Kaunergrathütte for their hospitality and technical support. Also, we would like to thank Eva Petrová and Dana Švehlová for assisting in molecular methods and analyses. Additionally, we are grateful to Dr. Eva Kaštovská for use of her lab and assistance.

## Author Contributions

KŘ, RA, JD, ATR, and JS designed the study. KŘ, RA, JD, ATR, JS, KČ, ZC, and RCM performed the field work. ATR, KČ, KŘ performed the lab work. ATR performed the data analysis. ATR wrote the manuscript with significant contributions from JS, JD, KŘ, RA, ZC and NP. TBM and PI were consulted and advised interpretations of results. All coauthors read and approved the final version of this manuscript

## Open Research Statement

The short-read amplicon sequencing data have been deposited under the NCBI BioProject with accessions XXX (will be submitted upon publication). For reproducibility, reusability, and transparency, the scripts and data used in this study were deposited to GitHub and are available via Zenodo (REF) (will be submitted upon publication). All functions and packages are cited within the paper without any associated novel code being produced for the results.

## Abbreviations

NSC: Non-structural Carbohydrate
m.a.s.l.: meters above sea level
MBC: Microbial biomass carbon
MBN: Microbial biomass nitrogen
°C: degrees Celsius
OM: organic matter (%)
TN: total nitrogen
NH-_4_^+^-N: ammonium
TP: total phosphorus

## Supplemental Figures and Tables

**Supplementary Table 1.** Results of linear mixed effect models of all measured parameters including plant nutrient concentrations, rhizosphere soil measurements, soil biological parameters, and alpha diversity metrics of bacterial and fungal communities. Location was used as a covariate to account for repeated seasonal sampling and p-values were adjusted using False-Discovery Rate (FDR) correction.

| Variable |  | All Locations |  |  |  |  | Alpine Meadow |  |  | Subnival Zone |  |  |
| --- | --- | --- | --- | --- | --- | --- | --- | --- | --- | --- | --- | --- |
|  |  | Season | Plant | Habitat | Plant<br>x<br>Season | Habitat<br>x<br>Season | Season | Plant | Plant<br>x<br>Season | Season | Plant | Plant<br>x<br>Season |
| Plant Parameters | Leaf N (g g<sup>-1</sup>) | <0.001 | <0.001 | 0.016 | <0.001 | 0.263 | <0.001 | <0.001 | 0.492 | <0.001 | <0.001 | <0.001 |
|  | Leaf C (g g<sup>-1</sup>) | 0.040 | 0.139 | 0.567 | <0.001 | 0.645 | 0.013 | 0.035 | 0.371 | <0.001 | 0.563 | <0.001 |
|  | δ<sup>15</sup>N ‰ | 0.008 | <0.001 | 0.034 | 0.003 | 0.018 | 0.018 | <0.001 | 0.034 | 0.044 | <0.001 | 0.034 |
|  | δ<sup>13</sup>C ‰ | 0.063 | <0.001 | 0.132 | 0.010 | 0.030 | 0.125 | 0.064 | 0.821 | <0.001 | <0.001 | <0.001 |
|  | Leaf C:N (mol) | 0.002 | <0.001 | 0.069 | 0.003 | 0.148 | 0.001 | <0.001 | 0.609 | <0.001 | <0.001 | <0.001 |
|  | Root P (g g<sup>-1</sup>) | 0.002 | 0.001 | 0.110 | 0.133 | 0.809 | 0.011 | 0.642 | 0.482 | 0.052 | <0.001 | 0.052 |
|  | Root C (g g<sup>-1</sup>) | 0.013 | 0.011 | 0.895 | <0.001 | 0.264 | 0.026 | 0.011 | 0.008 | <0.001 | 0.130 | 0.012 |
|  | Root N (g g<sup>-1</sup>) | 0.103 | <0.001 | 0.756 | 0.004 | 0.756 | 0.110 | <0.001 | 0.110 | 0.030 | 0.006 | 0.004 |
|  | Root C:N (mol) | 0.884 | <0.001 | 0.884 | <0.001 | 0.884 | 0.210 | <0.001 | 0.002 | 0.116 | <0.001 | 0.017 |
|  | Root Starch (%) | 0.999 | 0.999 | 0.228 | <0.001 | 0.049 | 0.999 | 0.840 | 0.004 | 0.999 | 0.999 | <0.001 |
|  | Root Fructan (%) | 0.226 | <0.001 | 0.795 | 0.009 | 0.226 | 0.228 | 0.226 | 0.226 | 0.006 | <0.001 | 0.016 |
|  | Root Glucose (%) | 0.755 | <0.001 | 0.5539 | <0.001 | <0.001 | 0.755 | <0.001 | 0.554 | 0.755 | <0.001 | <0.001 |
|  | Root Sucrose (%) | 0.098 | <0.001 | 0.6535 | <0.001 | <0.001 | 0.042 | <0.001 | <0.001 | 0.7668 | <0.001 | 0.6586 |
|  | Root Fructose (%) | **0.014** | **<0.001** | 0.754 | **<0.001** | **0.008** | **0.028** | **<0.001** | 0.051 | **<0.001** | **<0.001** | **<0.001** |
| Soil | Extractable Organic C (ug g<sup>-1</sup>) | **<0.001** | **0.016** | **0.001** | **0.048** | 0.176 | **<0.001** | 0.207 | 0.075 | **0.048** | **0.001** | 0.096 |
|  | Extractable N (ug g<sup>-1</sup>) | **<0.001** | **0.002** | **0.001** | **0.003** | 0.067 | **<0.001** | 0.067 | **0.038** | 0.088 | **0.001** | **0.004** |
|  | Extractable C:N (mol) | 0.084 | **0.002** | 0.763 | **<0.001** | 0.186 | 0.063 | **0.006** | **<0.001** | 0.104 | **0.046** | 0.186 |
|  | Rhizosphere pH | 0.127 | **<0.001** | 0.786 | **<0.001** | 0.321 | 0.209 | **<0.001** | **0.006** | 0.212 | **<0.001** | **0.008** |
| Biological Soil Parameters | Betaglucosidase | **<0.001** | **0.021** | 0.056 | 0.056 | 0.124 | **<0.001** | **<0.001** | **<0.001** | 0.997 | 0.969 | 0.969 |
|  | Cellobiosidase | **<0.001** | **0.018** | 0.075 | 0.054 | 0.166 | **<0.001** | **<0.001** | **<0.001** | 0.979 | 0.979 | 0.979 |
|  | Phosphatase | **<0.001** | **0.017** | **0.012** | 0.131 | 0.204 | **<0.001** | **0.002** | **0.011** | 0.982 | 0.417 | 0.982 |
|  | Leucinaminopeptidase | 0.070 | 0.168 | 0.070 | 0.141 | 0.112 | 0.070 | 0.070 | 0.070 | 0.887 | 0.910 | 0.620 |
|  | Chitinase | **<0.001** | 0.063 | **0.004** | 0.227 | 0.112 | **<0.001** | **<0.001** | **0.002** | 0.988 | 0.917 | 0.988 |
|  | MBC (ug g<sup>-1</sup>) | **<0.001** | **<0.001** | **<0.001** | **<0.001** | 0.034 | **<0.001** | **<0.001** | **<0.001** | 0.333 | **0.009** | **0.034** |
|  | MBN (ug g<sup>-1</sup>) | **<0.001** | **<0.001** | **0.001** | **<0.001** | **0.010** | **<0.001** | **<0.001** | **<0.001** | 0.156 | **0.016** | **0.013** |
|  | Microbial C:N<sub>mol</sub> | 0.958 | 0.958 | 0.958 | 0.958 | 0.958 | 0.958 | 0.958 | 0.958 | 0.958 | 0.958 | 0.958 |
| Microbial Alpha Diversity | Bacterial ASV Richness | **<0.001** | 0.163 | 0.721 | **0.033** | 0.944 | **<0.001** | 0.163 | **0.012** | **0.004** | 0.352 | 0.721 |
|  | Bacterial H' | **0.007** | 0.834 | 0.834 | 0.834 | 0.907 | **0.007** | 0.823 | 0.823 | **0.007** | 0.870 | 0.834 |
|  | Bacterial Simpson | 0.713 | 0.999 | 0.999 | 0.999 | 0.999 | 0.388 | 0.713 | 0.999 | 0.226 | 0.999 | 0.999 |
|  | Fungal OTU Richness | 0.081 | **0.037** | **0.039** | 0.054 | 0.341 | 0.188 | 0.054 | 0.0607 | 0.177 | 0.061 | 0.177 |
|  | Fungal H' | 0.222 | 0.268 | 0.359 | 0.201 | 0.640 | 0.201 | 0.420 | 0.201 | 0.359 | 0.268 | 0.268 |
|  | Fungal Simpson | 0.444 | 0.444 | 0.780 | 0.322 | 0.772 | 0.081 | 0.810 | 0.333 | 0.272 | 0.444 | 0.333 |

**Supplementary Table 2.** Permanova results of microbial assemblages in each season and habitat assessing the effect of bulk soil vs. rhizosphere soil and distinct plant species. Significance is indicated by asterisks (* p < 0.05, ** <0.01).

| Season | Alpine Meadow |  |  |  |  |  | Subnival Zone |  |  |  |  |  |
| --- | --- | --- | --- | --- | --- | --- | --- | --- | --- | --- | --- | --- |
|  | Fungi |  |  | Bacteria |  |  | Fungi |  |  | Bacteria |  |  |
|  | TSS | Rhizo | Plant Species | TSS | Rhizo | Plant Species | TSS | Rhizo | Plant Species | TSS | Rhizo | Plant Species |
| Spring | 23.3<br>(n=27) | **6.9%**<br>** | **19.5**<br>** | 6.74<br>(n=22) | **9.5%**<br>** | **22.3%**<br>** | 22.9<br>(n=27) | **5.9%**<br>** | **20.0%**<br>** | 7.9<br>(n=27) | **11.8%**<br>** | **19.7**<br>** |
| Summer | 23.2<br>(n=27) | **5.6%**<br>** | **18.8%**<br>** | 6.69<br>(n=20) | **8.6%**<br>** | **26.8%**<br>** | 22.6<br>(n=28) | **6.5%**<br>** | **21.8%**<br>* | 7.9<br>(n=25) | **12.1%**<br>** | **21.0%**<br>** |
| Autumn | 20.7<br>(n=24) | **4.7%**<br>* | **18.7%**<br>** | 7.6<br>(n=23) | **6.3%**<br>* | **21.8%**<br>** | 24.9<br>(n=30) | **6.8%**<br>** | **20.4%**<br>** | 8.71<br>(n=28) | **6.9%**<br>** | **21.6%**<br>** |
| Winter | 6.20<br>(n=9) | **15.6%**<br>* | **15.7%**<br>* | 1.9<br>(n=9) | **39.8%**<br>** | 9.2% | 7.6<br>(n=10) | 9.8% | 11.6% | 2.91<br>(n=10) | **25.7%**<br>* | 11.7% |

**Supplementary Table 3.**
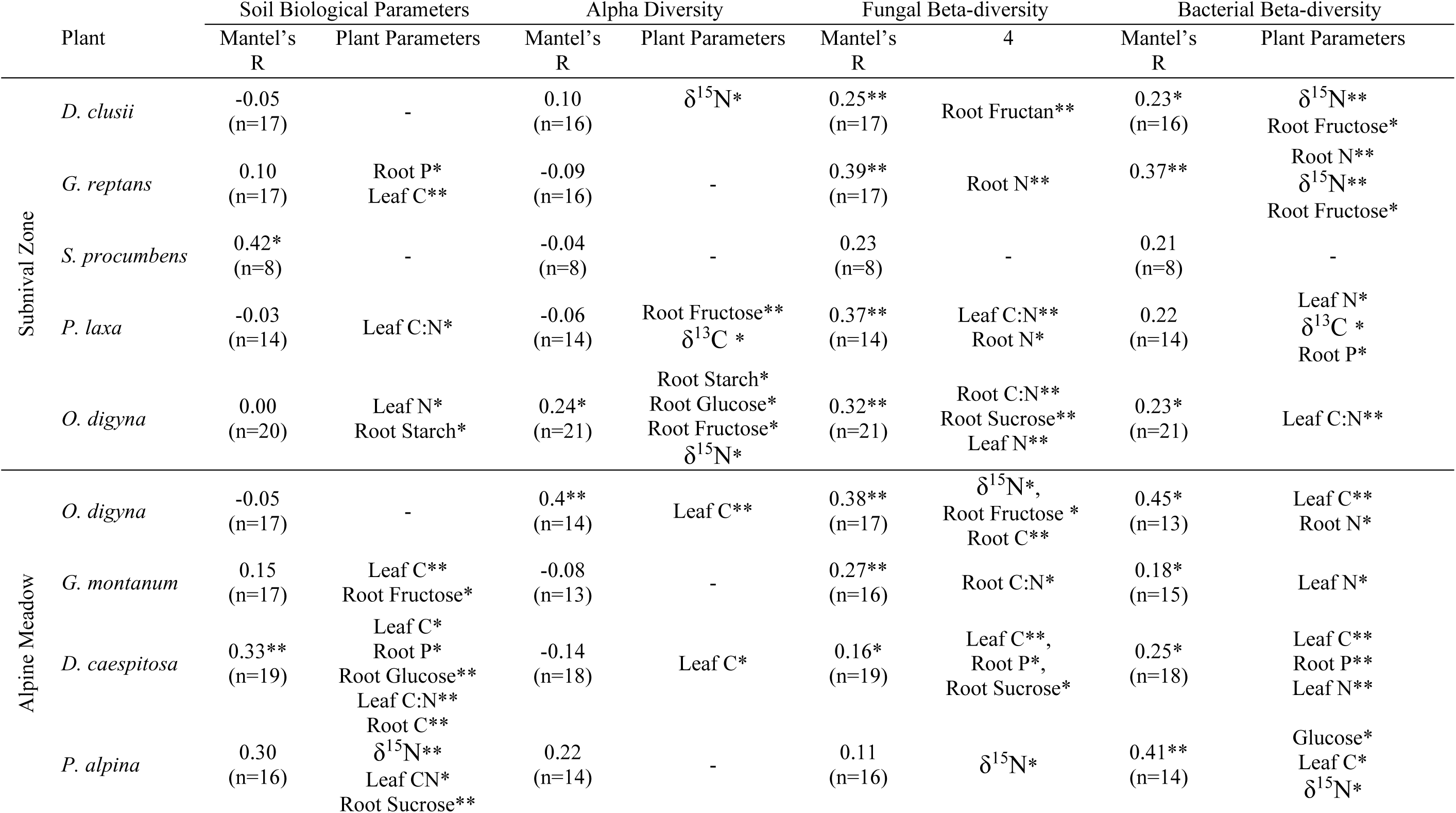
Mantel test results and forward-selected plant parameters across different levels of soil microbial measurements and different plant species. Significance is indicated by asterisks (* p < 0.05, ** <0.01). Soil biological parameters include microbial biomass carbon and nitrogen, and enzymatic potential. Alpha diversity is both bacterial and fungal H’ index, Simpson index, and richness.

**Supplementary Figure 1.**
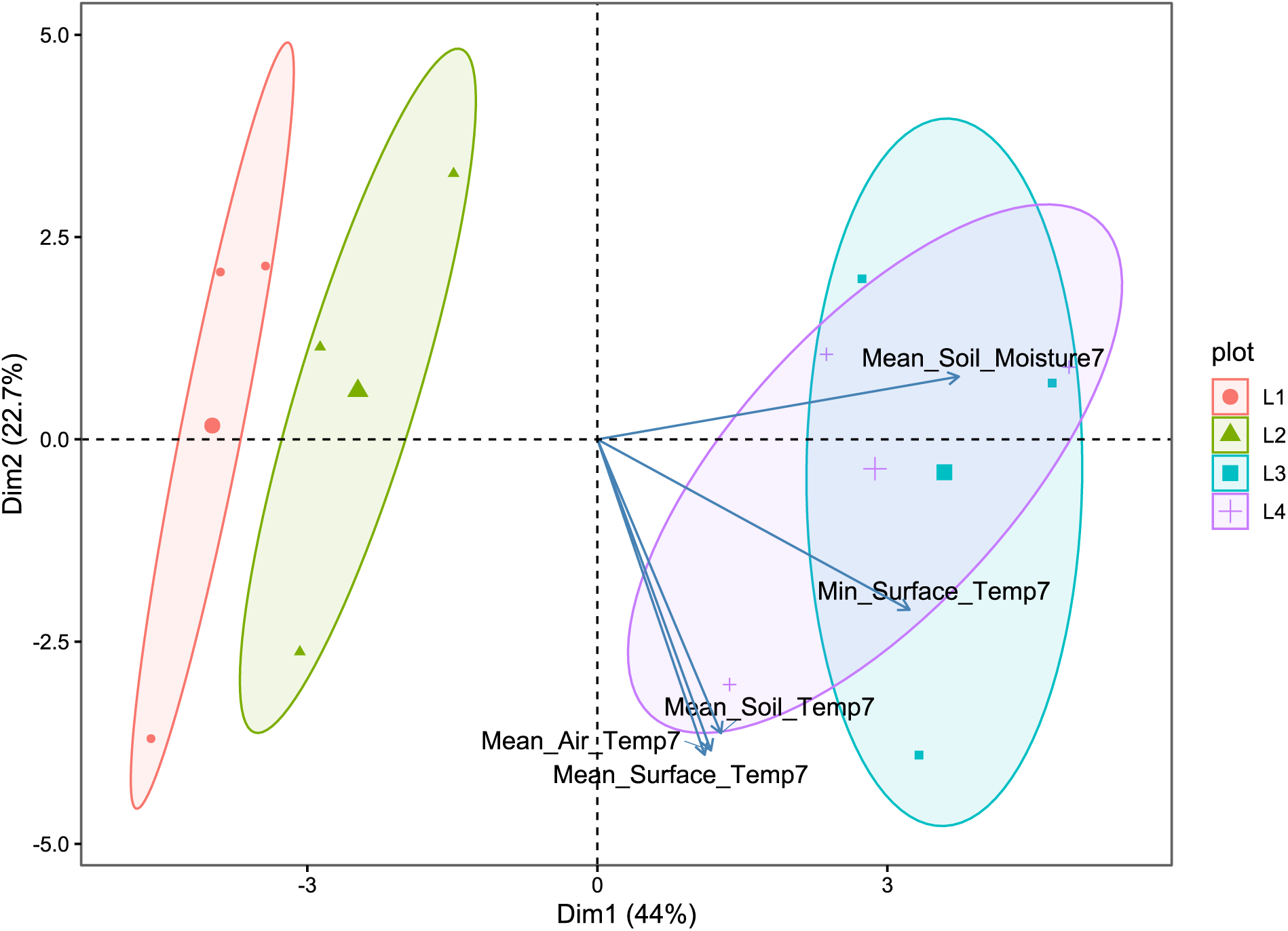
Principal component analysis of microclimatic parameters (air, surface, soil temperature, and soil moisture) and soil physicochemistry measurements collected during the growing season (spring, summer, autumn) at the four study locations.

**Supplementary Figure 2.**
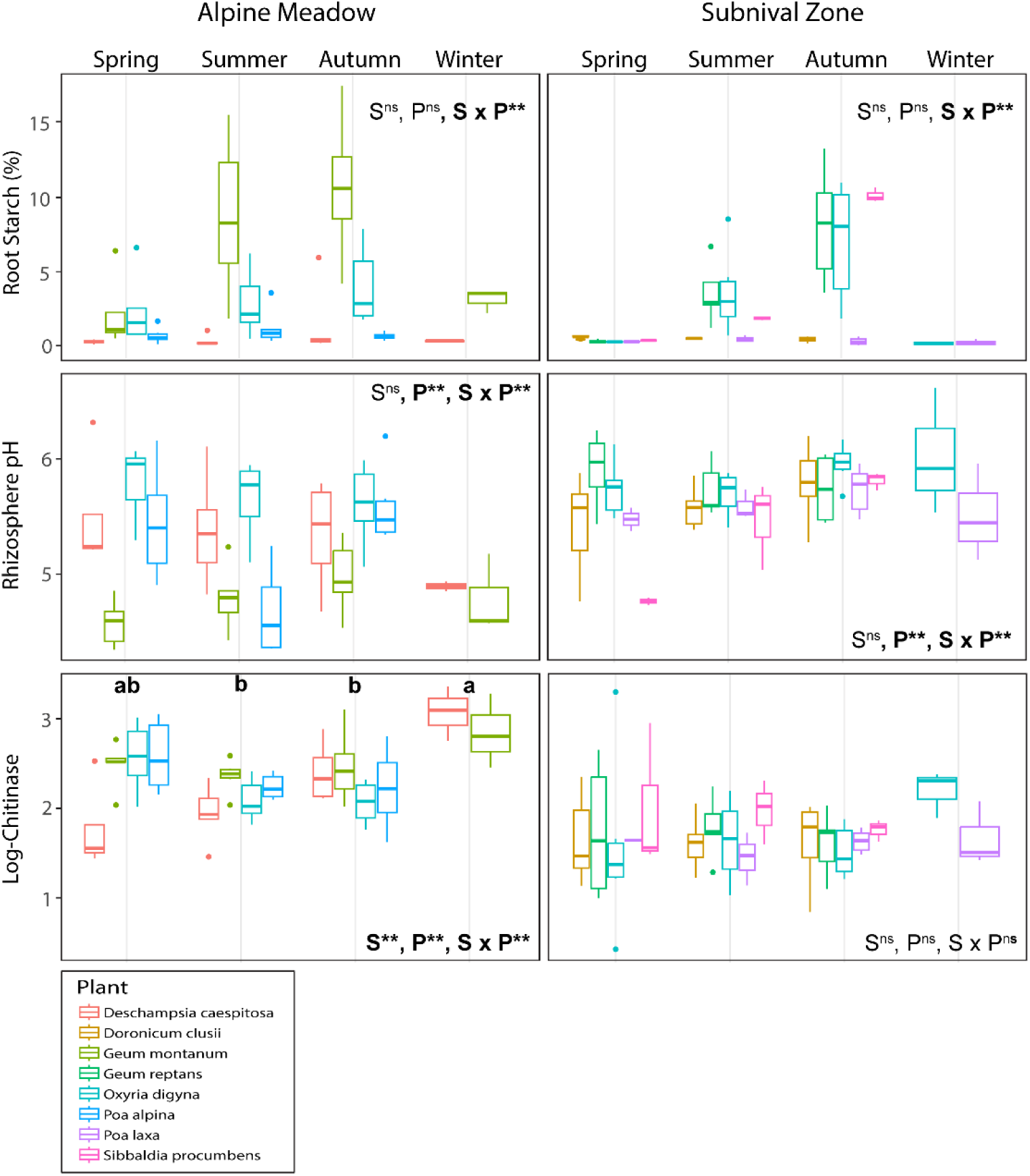
Box plots including linear mixed-effect model results testing the effect of season (S) and plant species (P) individually and their interaction (S x P) upon measured plant parameters within the rhizosphere of 8 different plant species occurring in the alpine meadow and subnival zone. Post-hoc Wilcoxon rank sum tests indicate seasonal differences amongst all plant species.

### Supplementary Plate A. *Doronicum clusii* (All.) Tausch

Clusius’ Leopard’s-bane

A geophyte (Asteraceae) flowering from July to August in central and southern Europe. It is characterized by rhizomatous growth, where an underground stem produces adventitious roots and stems annually. This species grows in rubble on silicate rock, primarily in subalpine to alpine regions.

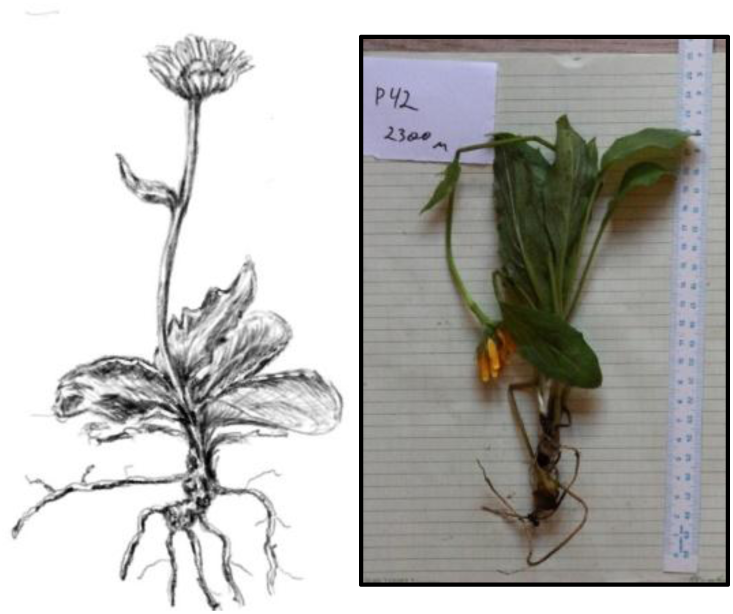

**Table 1.** Results of spearman rank correlation tests using forward-selected plant parameters as explanatory variables and rhizosphere parameters as response variables. Bold rows are visualized in individual plant phenologies (Figures 6 & 7).

| Exp. Var. | Resp. Var. | Spearman Corr. | P-value |
| --- | --- | --- | --- |
| $\delta^{15}\text{N}$ | **Bacterial Simpson Index** | **0.6784402** | **0.004** |
| **Fructan %** | ***Ascomycota inc. sed. sp.1*** | **-0.7113273** | **0.001** |
| $\delta^{15}\text{N}$ | Bacterial H' Index | 0.6107434 | 0.012 |
| $\delta^{15}\text{N}$ | Bacterial Richness | 0.5474615 | 0.028 |
| Fructose % | *Pseudomonas syringae* | 0.5147059 | 0.044 |

### Supplementary Plate B. *Geum reptans* L.

Creeping Avens

An herbaceous, rhizomatous perennial hemicryptophyte (Rosaceae), characterized by epigeal stolons. Flowering from July to August, it is typically found in rock debris and alluvium within alpine regions of central and southern Europe. Popular medicine attributes healing properties to all species of the genus *Geum*, due to the presence of the glycoside gein, particularly concentrated in the roots and leaves.

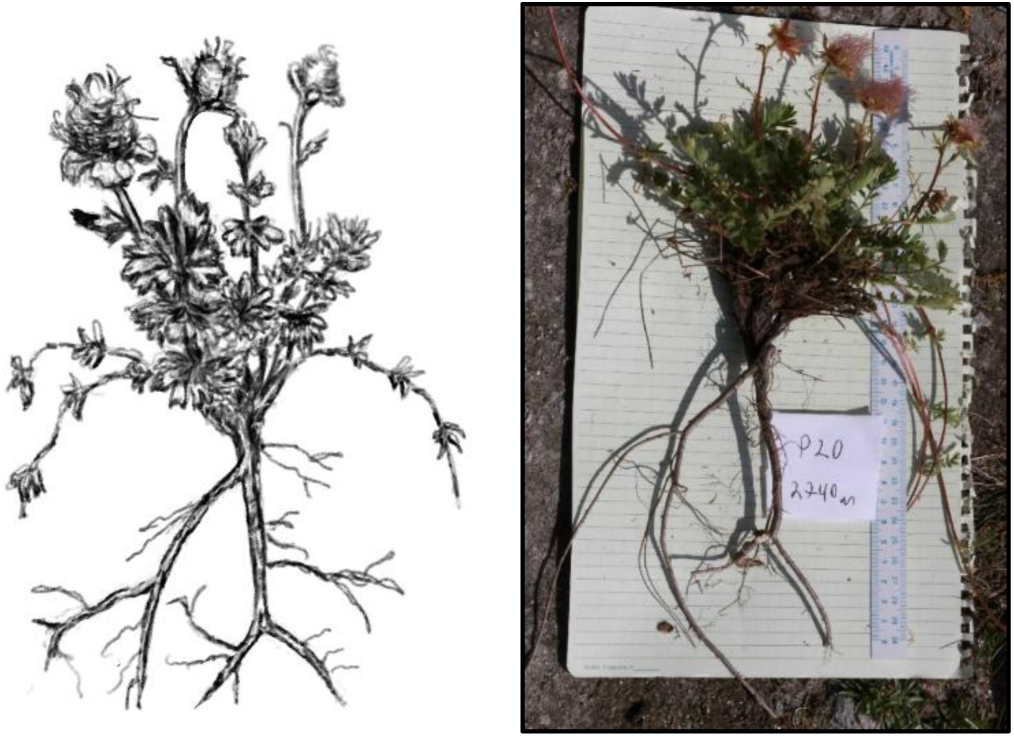

**Table 1.** Results of spearman rank correlation tests using forward-selected plant parameters as explanatory variables and rhizosphere parameters as response variables. Bold rows are visualized in individual plant phenologies (Figures 6 & 7).

| Exp. Var. | Resp. Var. | Spearman Corr. | P-value |
| --- | --- | --- | --- |
| **Leaf C** | **Microbial Biomass Carbon** | **0.71** | **0.001** |
| **Root P** | **Microbial Biomass Nitrogen** | **-0.67** | **0.003** |
| **Fructose %** | ***Serratia* sp. 1** | **0.66** | **0.005** |
| **Root P** | **Log-leucinaminopeptidase activity** | **-0.62** | **0.008** |
| Leaf C | Microbial Biomass Nitrogen | 0.60 | 0.011 |
| Leaf C | Log-betaglucosidase activity | 0.58 | 0.014 |
| Root N | *Herpotrichia pinotorum* | 0.54 | 0.026 |

### Supplementary Plate C. *Sibbaldia procumbens* L.

Creeping Sibbaldia

A perennial, herbaceous plant (Rosaceae) with a creeping growth habit and an oblique rhizome. Classified as both a subfructicose chamaephyte and a perennial hemicryptophyte, it flowers from June to July. It has an Arctic-alpine distribution within snow valleys of montane to alpine regions while also being grown as within lawns and flower beds.

No soil microbial variables were significantly correlated to above ground plant traits due to a limited sample size (n = 9).

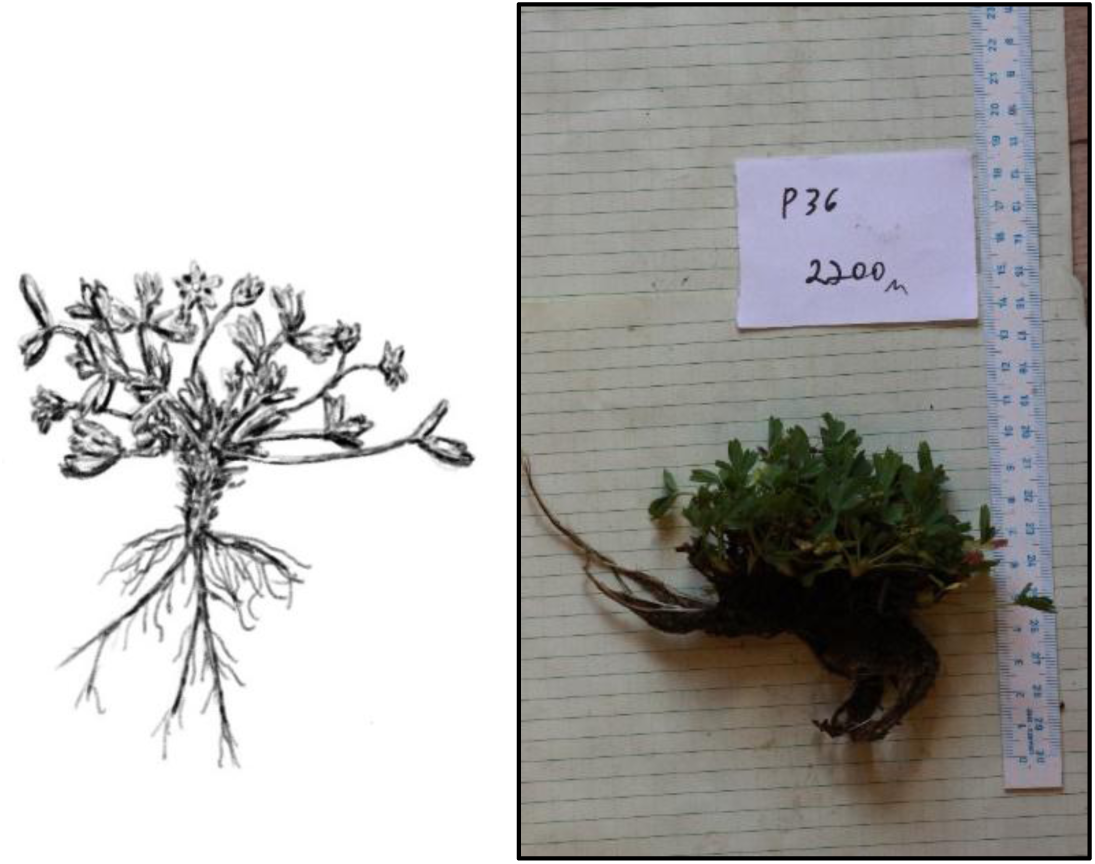

### Supplementary Plate D. *Poa laxa* Haenke

Limp Meadow-grass

A perennial hemicryptophyte (Poaceae) which grows in lime-free rock debris from subalpine to alpine regions. Found in central and southern Europe, it flowers from July to August.

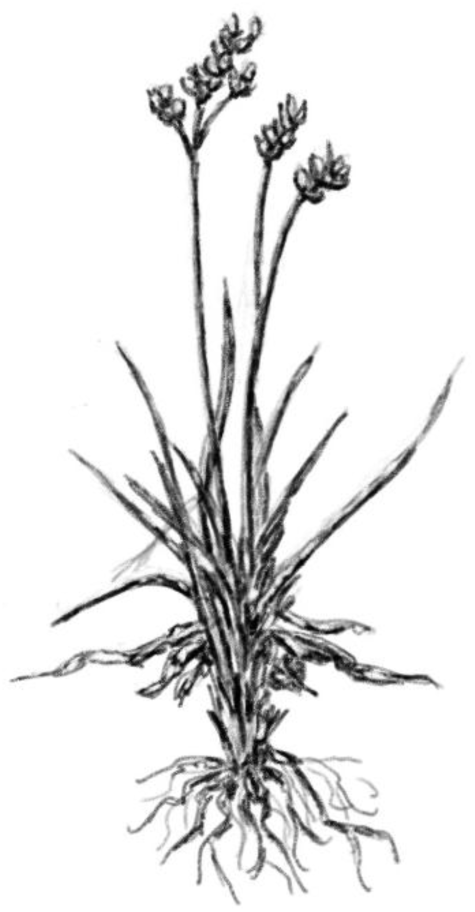

**Table 1.** Results of spearman rank correlation tests using forward-selected plant parameters as explanatory variables and rhizosphere parameters as response variables. Bold rows are visualized in individual plant phenologies (Figures 6 & 7).

| Exp. Var. | Res. Var. | Spearman Corr. | P-value |
| --- | --- | --- | --- |
| **Leaf C:N** | **Microbial Biomass Carbon** | **0.82** | **<0.001** |
| **Leaf C:N** | **Microbial Biomass Nitrogen** | **0.76** | **0.002** |
| **Leaf C:N** | **Helotiales *inc. sed.* Sp. 1** | **0.70** | **0.005** |
| **$\delta^{13}\text{C}$** | **Ktedonobacteraceae sp.1** | **0.68** | **0.007** |
| Leaf N | *Bradyrhizobium japonicum* | -0.59 | 0.026 |
| Root N | *Lachnum* sp. 1 | -0.59 | 0.027 |
| $\delta^{13}\text{C}$ | Bacterial H' Index | -0.56 | 0.037 |
| Leaf C:N | *Lachnum* sp. 1 | 0.55 | 0.04 |
| $\delta^{13}\text{C}$ | Fungal Simpson Index | 0.54 | 0.047 |

### Supplementary Plate E. *Oxyria digyna* (L.) Hill – Subnival

Mountain Sorrel

A hemicryptophyte scapose plant with leaves that contain oxalic acid and flowers from July to September. It grows in wet debris of rocks, siliceous screes, and moraine environments in the Alpine biome, often found on lime-poor, moist rock scree, and snow valleys with flowing water. The deep tap-root helps to anchor the plant in long snow-covered terrain while also storing non-structural carbohydrates such as starch and fructan.

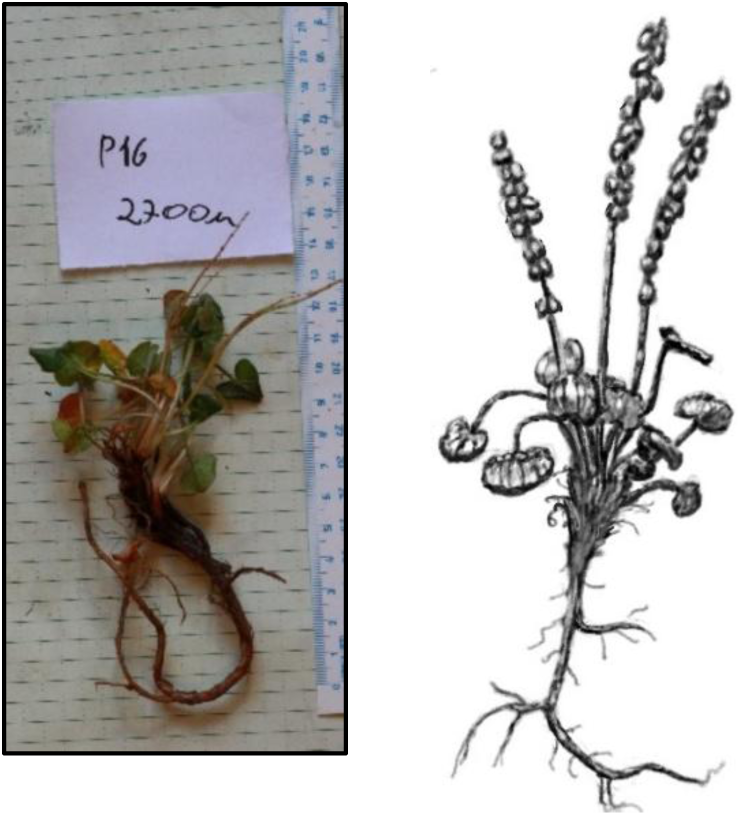

**Table 1.**
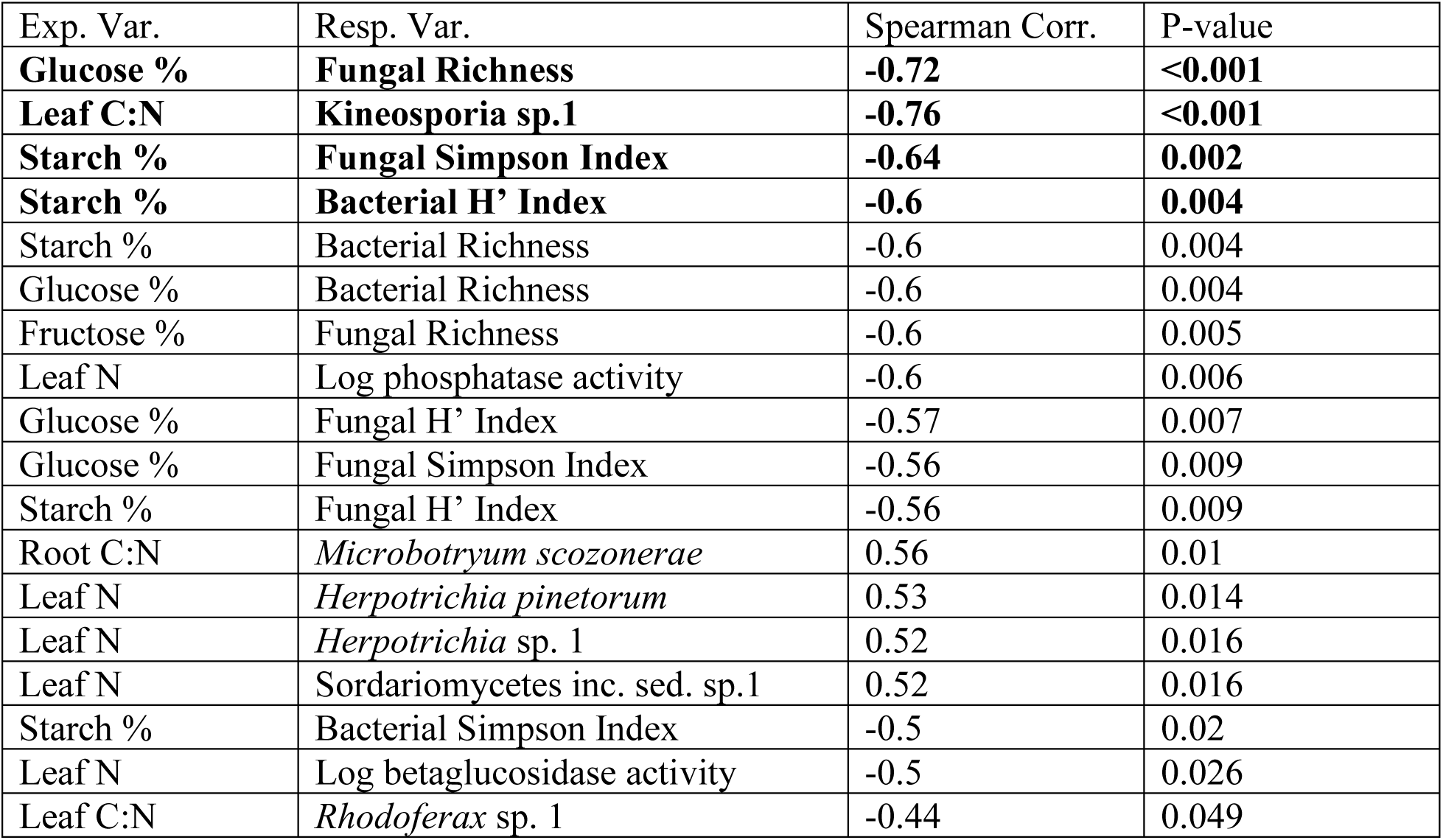
Results of spearman rank correlation tests using forward-selected plant parameters as explanatory variables and rhizosphere parameters as response variables. Bold rows are visualized in individual plant phenologies (Figures 6 & 7).

### Supplemental Plate F. *Oxyria digyna* (L.) Hill - Alpine Meadow

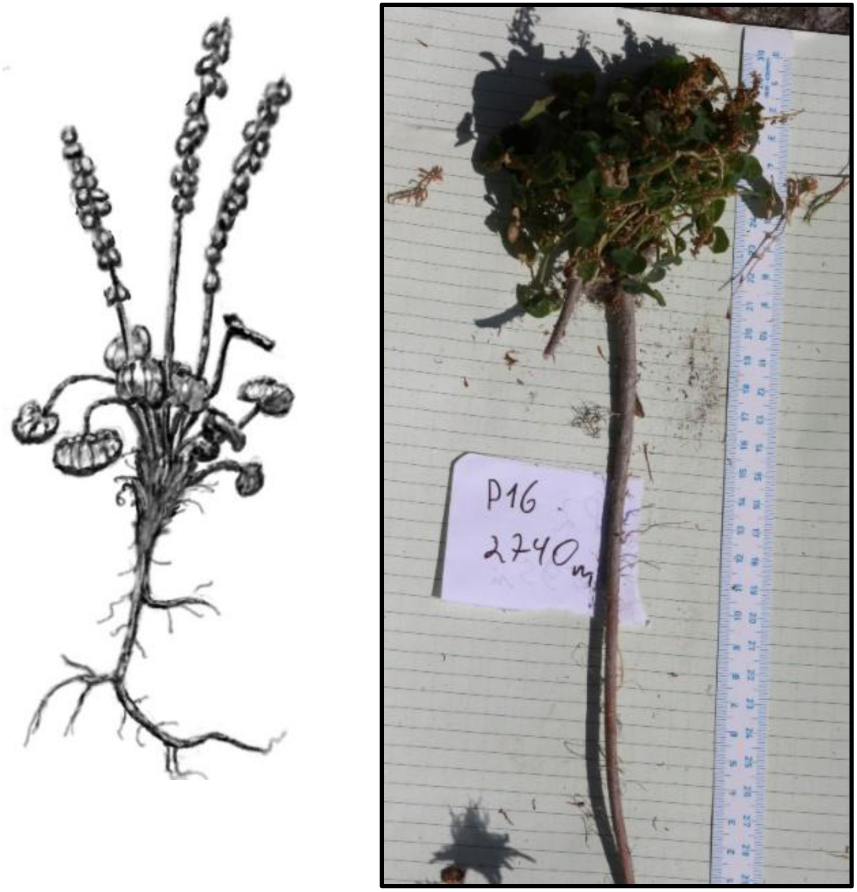

**Table 1.** Results of spearman rank correlation tests using forward-selected plant parameters as explanatory variables and rhizosphere parameters as response variables. Bold rows are visualized in individual plant phenologies (Figures 6 & 7).

| Exp. Var. | Resp. Var. | Spearman Corr. | P-value |
| --- | --- | --- | --- |
| **Root C** | ***Pezicula heterochroma*** | **0.75** | **0.001** |
| **Fructose %** | ***Pezicula heterochroma*** | **0.71** | **0.001** |
| **Root N** | ***Serratia sp. 1*** | **0.72** | **0.006** |
| **Root C** | ***Microbotryum scozoneræ*** | **0.62** | **0.008** |
| **Leaf C** | **Fungal Simpson Index** | **0.72** | **0.011** |
| Leaf C | Fungal OTU Richness | 0.69 | 0.014 |
| Root N | Comamodaceae sp.1 | -0.65 | 0.016 |
| Leaf C | Fungal H' Index | 0.66 | 0.024 |
| Leaf C | Bacterial ASV Richness | 0.67 | 0.032 |
| Fructose % | *Microbotryum scozoneræ* | 0.5 | 0.04 |

### Supplementary Plate G. *Geum montanum* L.

Alpine Avens

A clonal, perennial hemicryptophyte (Rosaceae) with buds at ground level and leaves arranged in a basal rosette, growing from a horizontal rhizome. It grows in grasslands and pastures from montane to subalpine and alpine regions of central and southern Europe, flowering from May to August.

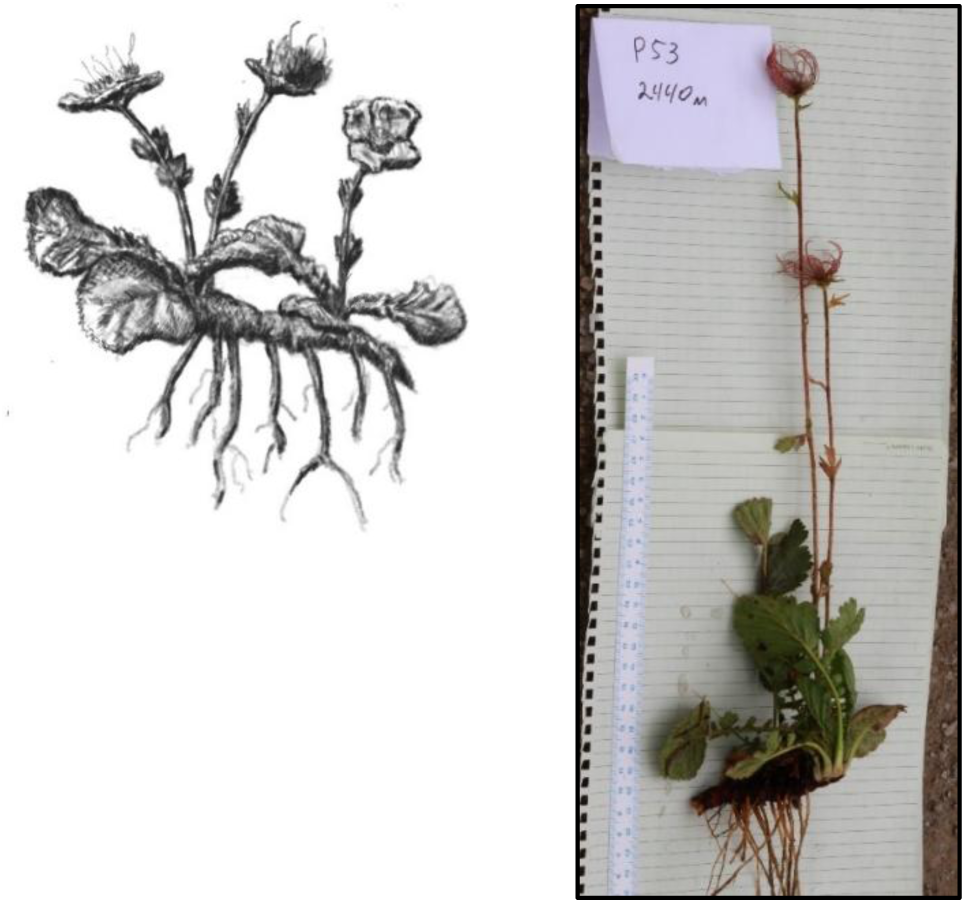

**Table 1.** Results of spearman rank correlation tests using forward-selected plant parameters as explanatory variables and rhizosphere parameters as response variables. Bold rows are visualized in individual plant phenologies (Figures 6 & 7).

| Exp. Var. | Res. Var. | Spearman Corr. | P-value |
| --- | --- | --- | --- |
| **Fructose %** | **Microbial Biomass C:N** | **0.64** | **0.003** |
| **Leaf C** | **Log Betaglucosidase activity** | **0.6** | **0.005** |
| **Root C:N** | ***Cladophialophora* sp.1** | **-0.66** | **0.005** |
| Leaf C | Log Cellobiosidase activity | 0.55 | 0.012 |
| Leaf C | Log Phosphatase activity | 0.52 | 0.018 |
| Leaf N | *Rhizorhapis* sp.1 | 0.59 | 0.019 |
| Leaf N | *Kineosporia* sp.1 | 0.6 | 0.02 |
| Leaf C | Log Chitinase activity | 0.5 | 0.024 |
| Fructose % | Log Leucinaminopeptidase activity | -0.47 | 0.04 |

### Supplementary Plate H. *Deschampsia cespitosa* (L.) P. Beauv.

Tufted Hairgrass

A cespitose hemicryptophyte (Poaceae), that thrives in wet soils, along roadsides, banks, and in forests to alpine regions, blooming from June to August. Highly tolerant and resilient species with a wordwide distribution.

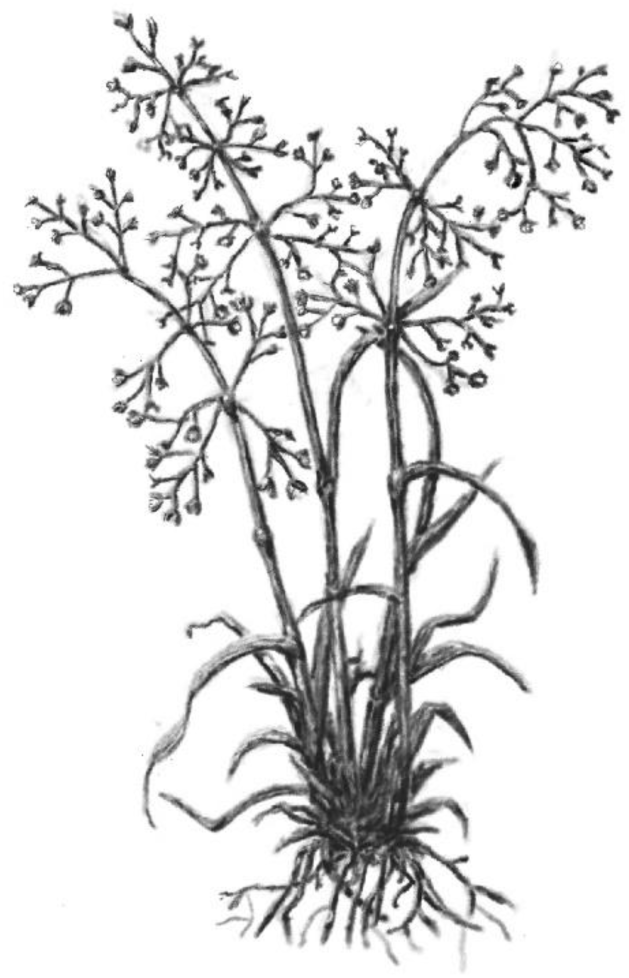

**Table 1.** Results of spearman rank correlation tests using forward-selected plant parameters as explanatory variables and rhizosphere parameters as response variables. Bold rows are visualized in individual plant phenologies (Figures 6 & 7).

| Exp. Var | Resp. Var. | Spearman Corr. | P-value |
| --- | --- | --- | --- |
| **Leaf C** | **Microbial Biomass Nitrogen** | **0.72** | **<0.001** |
| **Glucose %** | **Log Leucinaminopeptidase Activity** | **-0.7** | **<0.001** |
| **Leaf C** | **Log Phosphatase activity** | **0.69** | **0.001** |
| **Glucose %** | **Microbial Biomass C:N** | **0.66** | **0.002** |
| **Leaf C** | **Microbial Biomass Carbon** | **0.63** | **0.004** |
| **Leaf C** | **Fungal H' Index** | **0.62** | **0.006** |
| Leaf C | *Pseudomonas frederiksbergensis* | 0.62 | 0.007 |
| Leaf C | Fungal Richness | 0.55 | 0.017 |
| Leaf C | *Acinetobacter* sp. 1 | -0.56 | 0.017 |
| Leaf C | *Eocronartium* sp.1 | 0.52 | 0.024 |
| Root P | *Pseudomonas chlororaphis* | 0.52 | 0.027 |
| Leaf C | *Klebsiella* sp. 1 | -0.51 | 0.032 |

### Supplementary Plate I. *Poa alpina* L. Alpine

Meadow-grass

A grass species in the Poaceae family, which thrives in meadows and pastures of subalpine to alpine regions, flowering from June to September. Distributed across Eurosiberian and North American regions. This perennial grass prefers well-draining soils and full sun.

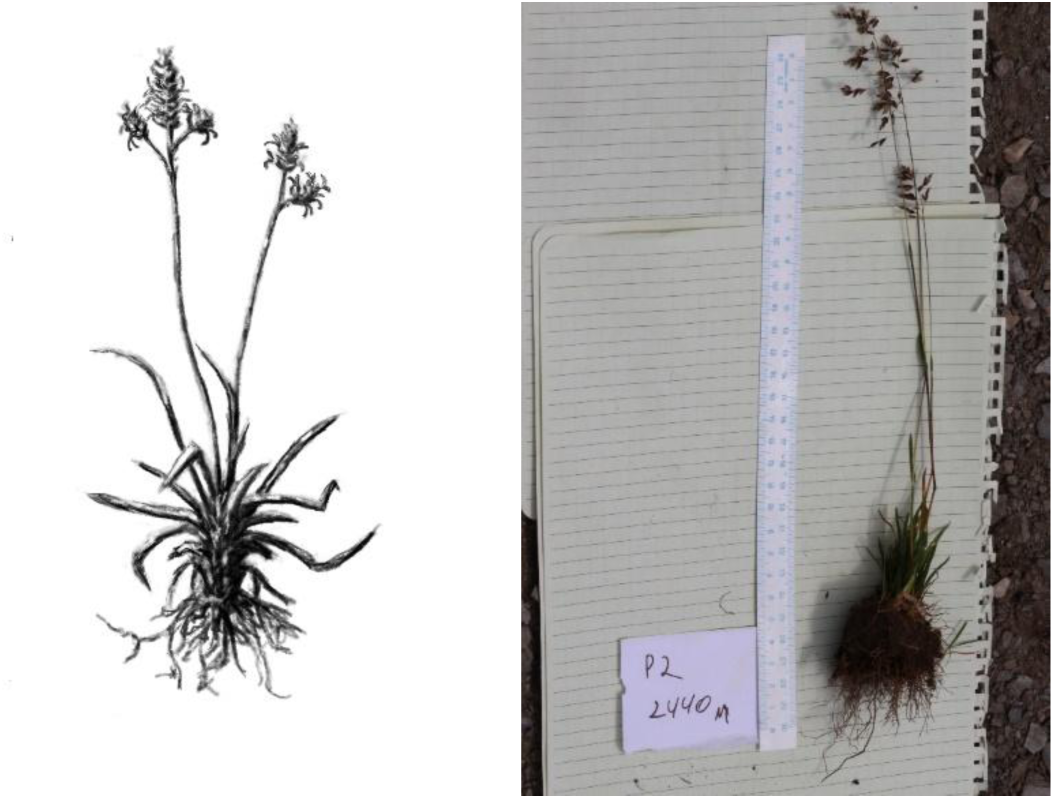

**Table 1.** Results of spearman rank correlation tests using forward-selected plant parameters as explanatory variables and rhizosphere parameters as response variables. Bold rows are visualized in individual plant phenologies (Figures 6 & 7).

| Exp. Var. | Resp. Var. | Spearman Corr. | P-value |
| --- | --- | --- | --- |
| $\delta^{15}\text{N}$ | *Burkholderia xenovorans* | **0.7734811** | **0.001** |
| **Root C** | **Microbial Biomass Carbon** | **0.7058824** | **0.003** |
| **Root C** | **Microbial Biomass Nitrogen** | **0.6470588** | **0.008** |
| Leaf C:N | Microbial Biomass C:N | 0.6323529 | 0.01 |
| Leaf C:N | Log Leucinaminopeptidase activity | -0.5529412 | 0.023 |
| $\delta^{15}\text{N}$ | Comamodaceae sp.1 | -0.5556811 | 0.039 |

